# A *Coxiella burnetii* effector interacts with the host PAF1 complex and suppresses the innate immune response

**DOI:** 10.1101/2022.04.20.488957

**Authors:** Natasha Lopes Fischer, Mark A. Boyer, William P. Bradley, Lynn A. Spruce, Hossein Fazelinia, Sunny Shin

## Abstract

Intracellular bacteria such as the pathogen *Coxiella burnetii* inject effector proteins into the host cell that promote productive infection. One common strategy of pathogen effectors is to suppress host immune responses to enable pathogen replication. The *C. burnetii* type IV secretion system translocates a large number of effectors into host cells that collectively promote intracellular bacterial replication, but the individual functions of most of these effectors are poorly understood. In this study, we describe a *C. burnetii* effector, CBU1314, that localizes to the nucleus and inhibits NF-κB-, MAPK-, and type I IFN-dependent gene expression. Mechanistically, we find that CBU1314 interacts with the PAF1 complex (PAF1C), a central host transcriptional complex that regulates expression of inflammatory genes in innate immune cells. Notably, we find that PAF1 promotes immune gene expression in response to various immune agonists and *C. burnetii* infection. Moreover, we find that PAF1 is critical for restricting intracellular *C. burnetii* replication. Overall, our findings uncover PAF1C as a host target of a *C. burnetii* effector and reveal new insight into how intracellular bacterial pathogens subvert cell-intrinsic innate defenses.

**Significance:** Intracellular bacteria often employ secreted effector proteins to modulate cellular processes and survive intracellularly. The study of these effectors can provide valuable insight into microbial pathogenesis and host biology. Here, we describe a *Coxiella burnetii* effector that inhibits multiple host signaling pathways and modulates the innate immune response. This effector interacts with the host PAF1 complex (PAF1C), a central regulator of transcription. Furthermore, we show that PAF1 is necessary for maximal gene expression downstream of various innate immune receptors and restricts bacterial replication during infection. Overall, our study elucidates the function of a *C. burnetii* effector in evading the immune response and provides insight into the role of PAF1C in cell-intrinsic defense against bacterial pathogens.

## Introduction

*Coxiella burnetii* is a highly infectious and underdiagnosed zoonotic gram-negative pathogen that causes the flu-like illness Q fever. Humans usually become infected with *C. burnetii* by breathing in aerosolized bacteria that are shed from livestock. Following inhalation, *C. burnetii* infects and replicates in alveolar macrophages in the lung. While most Q fever cases are asymptomatic or self-limiting, acute infection can progress to serious chronic disease and life-threatening endocarditis that is difficult to treat (Maurin & Raoult, 1999).The largest Q fever outbreak reported occurred in the Netherlands from 2007-2010 with over 4000 cases.

Individuals living as far as 5km from farms with infected animals became ill, demonstrating the far-reaching consequences of this pathogen (Schneeberger, Wintenberger, van der Hoek, & Stahl, 2014). Because of its aerosolized large-scale transmission, environmental stability, and low infectious dose, *C. burnetii* is classified as a Category B select agent for its threat to human health and potential to be developed as a bioweapon (Madariaga, Rezai, Trenholme, & Weinstein, 2003; Rotz, Khan, Lillibridge, Ostroff, & Hughes, 2002). Despite this potential threat, the near-obligate lifestyle of this organism and technical challenges associated with genetic manipulation of this pathogen (Beare, 2012; Omsland et al., 2009; Omsland & Heinzen, 2011) have hampered studies into the molecular mechanisms of how it interacts with the innate immune system and replicates within host cells to cause disease.

*C. burnetii* uses a Type IVB Secretion System (T4SS) to inject over 150 bacterial effector proteins into the host cell (Larson et al., 2016). These effectors are thought to modulate many cellular processes to allow the *Coxiella-*containing vacuole to mature into an acidic phagolysosome-like compartment where the bacteria can replicate (Moffatt, Newton, & Newton, 2015). While the functions of most effectors are still unknown, some effectors have been found to suppress innate immune responses. *C. burnetii* has four effectors (AnkG, CaeA, CaeB, IcaA) that suppress cell death pathways (Bisle et al., 2016; Cordsmeier et al., 2022; Cunha et al., 2015; Klingenbeck, Eckart, Berens, & Lührmann, 2013; Lührmann, Nogueira, Carey, & Roy, 2010), three effectors (CBU0885, CBU1676, CBU0388) that modulate the Cell Wall Integrity MAPK pathway in yeast (Lifshitz et al., 2014), and two recently discovered effectors (NopA and CinF) that suppress NF-κB signaling in human cells (Burette et al., 2020; Y. Zhang et al., 2022). For many years, *C. burnetii* was considered to be an obligate intracellular pathogen until the relatively recent development of an axenic culture system (Omsland et al., 2009). Due to its obligate intracellular nature and intimate association with host cells, it is not surprising that *C. burnetii* possesses several effectors that suppress host immune responses, and it is likely that more exist.

During infection, pattern recognition receptors (PRRs) at the cell surface and within the cell detect pathogen components and activities. PRRs then initiate signaling cascades that culminate in host gene expression and the upregulation of antimicrobial programs (Janeway & Medzhitov, 2002; Medzhitov, 2007). NF-κB and MAPK are central signaling pathways activated by many immune receptors, such as the Toll-like receptors (TLRs), TNF receptors, and IL-1 receptors, and are key mediators of host defense (Kawai & Akira, 2011). Furthermore, IFN signaling leads to potent induction of interferon-stimulated genes (ISGs) and antimicrobial programs (Medzhitov, 2007; Platanias, 2005). Many pathogens have evolved ways to suppress these signaling pathways to dampen the immune response during infection (Roy & Mocarski, 2007), including numerous effectors that are secreted by the virulence-associated secretion systems of gram-negative bacterial pathogens (Finlay & McFadden, 2006; Rahman & McFadden, 2011).

Given the large number of T4SS-translocated effectors possessed by *C. burnetii*, its near-obligate host-associated lifestyle, and its ability to infect a broad array of mammalian species (Larson et al., 2016; Maurin & Raoult, 1999; Omsland et al., 2009), we considered it likely that *C. burnetii* possesses multiple effectors that target innate immune signaling pathways. In this study, we demonstrate that *C. burnetii* uses its T4SS to dampen innate immune responses in both immortalized and primary human myeloid cells. We identify a *C. burnetii* T4SS-translocated effector (CBU1314), that associates with host chromatin and broadly suppresses NF-κB, MAPK, and type I IFN signaling, thereby modulating gene expression downstream of multiple host innate immune receptors. Using immunoprecipitation followed by mass spectrometry analysis, we identify an interaction between CBU1314 and the host PAF1 complex (PAF1C), which was previously found to promote inflammatory and anti-viral gene expression, in part through regulating transcriptional elongation (Hou et al., 2019; Kenaston, Pham, Petit, & Shah, 2022; Kim, Guermah, & Roeder, 2010; Marazzi et al., 2012; Yu et al., 2015). Intriguingly, both influenza and dengue viruses possess non-structural (NS) proteins that antagonize PAF1C to dampen the immune response (Marazzi et al., 2012; Petit et al., 2021; Shah et al., 2018). We find that knockdown of PAF1 leads to suppression of NF-κB and IFN-dependent immune gene expression, similar to the effects of CBU1314 on host cells, suggesting that CBU1314 antagonizes PAF1C to suppress the innate immune response in host cells. Finally, we show that PAF1 restricts intracellular bacterial replication within host cells, demonstrating for the first time that PAF1C contributes to cell-intrinsic immune defense against an intracellular bacterial pathogen. These findings reveal a common strategy employed by viral and bacterial pathogens to target the host PAF1 complex in order to suppress immune gene expression.

## Results

### *C. burnetii* T4SS activity suppresses innate immune cytokine production in cell lines and primary human macrophages

*C. burnetii* activates multiple receptors and signaling pathways in innate immune cells, resulting in the production and secretion of inflammatory cytokines during infection (Ammerdorffer et al., 2015; Boucherit et al., 2012; Bradley et al., 2016; Dragan, Kurten, & Voth, 2019; Graham et al., 2013; Graham, Winchell, Kurten, & Voth, 2016; Meghari et al., 2005; Ramstead et al., 2016; Voth & Heinzen, 2009; Zamboni et al., 2004). However, C*. burnetii* also uses its T4SS activity to suppress phosphorylation of the NF-κB protein RelA during infection of the human monocytic THP-1 cell line (Mahapatra et al., 2016), and effectors have been recently identified that inhibit NF-κB signaling and nuclear import of transcription factors associated with the immune response (Burette et al., 2020; Y. Zhang et al., 2022). *C. burnetii* T4SS activity suppresses the induction of several inflammatory cytokines at the mRNA level and the cytokines TNF and IFN- α4 at the protein level in THP-1 cells (Burette et al., 2020). We wanted to examine whether the production and secretion of additional cytokines were also suppressed in a T4SS-dependent manner by *C. burnetii* and whether this suppression extended to primary human macrophages. We first infected THP-1 monocytes with WT *C. burnetii* or an *icmL* mutant lacking a functional T4SS (ΔT4SS) that is incapable of injecting effectors, and analyzed the mRNA expression of proinflammatory cytokines. WT bacteria-infected cells exhibited decreased expression of these genes compared to infection with the ΔT4SS strain (Fig. 1A), in agreement with previous findings (Burette et al., 2020).

**Figure 1.**
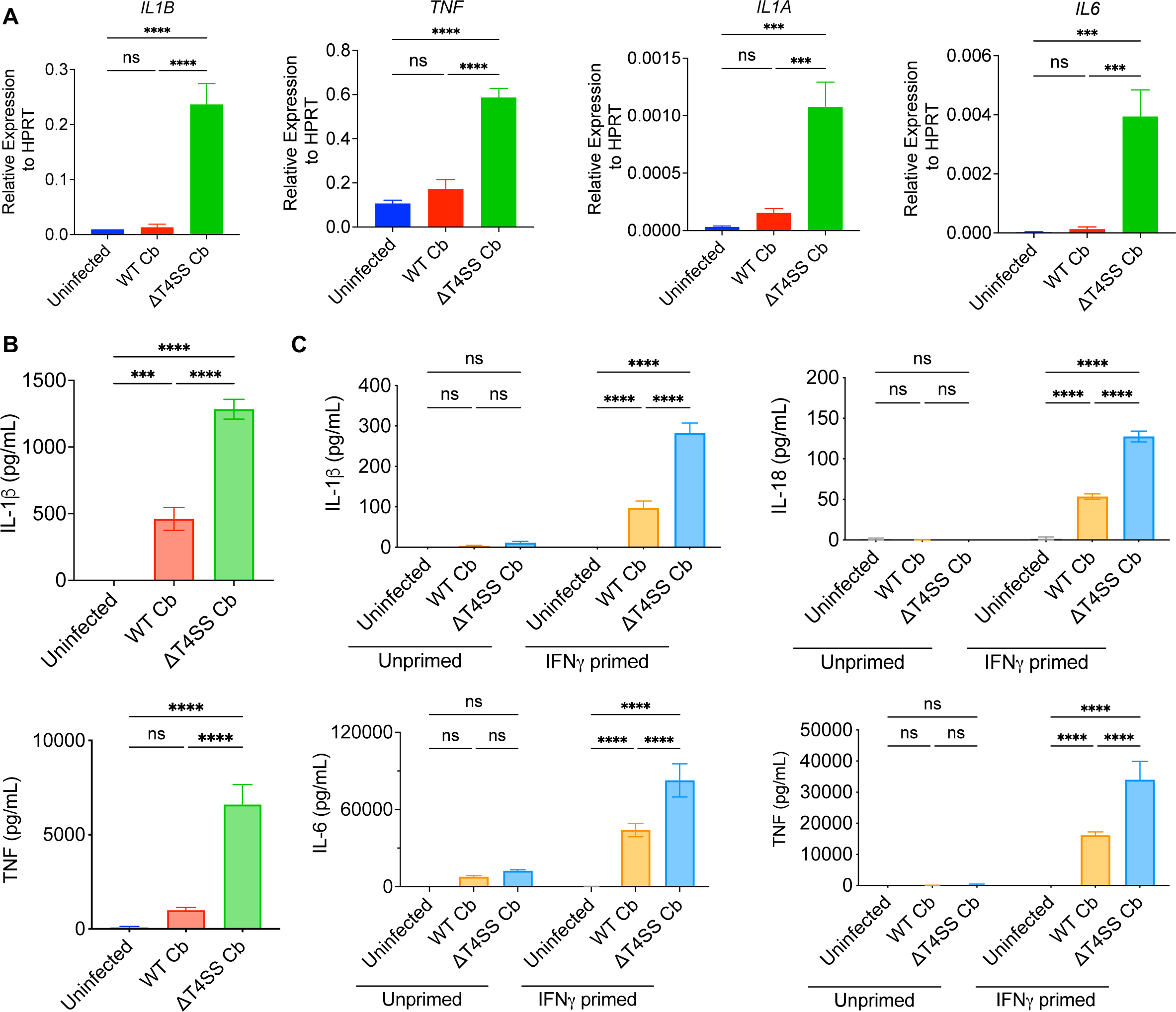
*C. burnetii* T4SS activity suppresses innate immune responses in THP-1 cells and primary human macrophages. A) THP-1 monocytes were infected with WT *C. burnetii* (WT Cb) or an *icmL*::*Tn* strain lacking a functional T4SS (ΔT4SS Cb), and mRNA expression was analyzed for the indicated genes at 24h.p.i. B) PMA-differentiated THP-1 macrophages were left uninfected or infected with WT or ΔT4SS *C. burnetii*. 24h.p.i. The supernatant was harvested and cytokine secretion was measured by ELISA. C) hMDMs were left unprimed or primed overnight with 100U/ml IFNγ, then infected and harvested as in B. Bar graphs display the mean +/- SD of triplicate replicates. Data is representative of 2-3 independent experiments for A and B, and three independent donors for C. Statistical analyses for A-B: One-way ANOVA with Tukey’s multiple comparisons test; for C: Two-way ANOVA with Tukey’s multiple comparisons test within each unprimed or primed group.

To examine cytokine secretion, we infected PMA-differentiated THP-1 macrophages. In line with the monocyte mRNA expression data, THP-1 macrophages secreted significantly less IL-1β and TNF in response to WT *C. burnetii* infection compared to infection with the ΔT4SS mutant (Fig. 1B). Importantly, we observed similar results in primary human monocyte-derived macrophages (hMDMs), whereby priming with interferon-gamma (IFN-γ) to boost pathogen detection (Kak, Raza, & Tiwari, 2018) revealed that WT *C. burnetii* also suppresses cytokine secretion in these cells in a T4SS-dependent manner (Fig. 1C). Collectively, these results indicate that *C. burnetii* T4SS activity suppresses innate immune cytokine production in both THP-1 cells and primary human macrophages.

### Identification of a *C. burnetii* effector that suppresses NF-κB signaling

Our data, along with previous findings (Burette et al., 2020), indicated that *C. burnetii* utilizes its T4SS to suppress innate cytokine expression. As many innate immune cytokines rely on NF-κB signaling for their expression (Kopp & Ghosh, 1995; Smale, 2012), we hypothesized that *C. burnetii* must utilize one or multiple effectors to suppress NF-κB signaling. We therefore conducted an NF-κB luciferase reporter screen using a subset of ectopically expressed *C. burnetii* effectors (Fig. 2A-B). Mammalian vectors encoding GFP-tagged effectors were transfected into HEK293T cells in conjunction with vectors encoding the *Firefly* luciferase gene driven by NF-κB-responsive promoter elements, and the *Renilla* luciferase gene driven by a constitutive actin promoter. Cells were stimulated with TNF to activate the NF-κB pathway, and luciferase activity was measured 4 hours after activation. Of the 35 effectors tested, the effector CBU1314 demonstrated the strongest suppression of NF-κB-induced luciferase expression (Fig. 2B).

**Figure 2.**
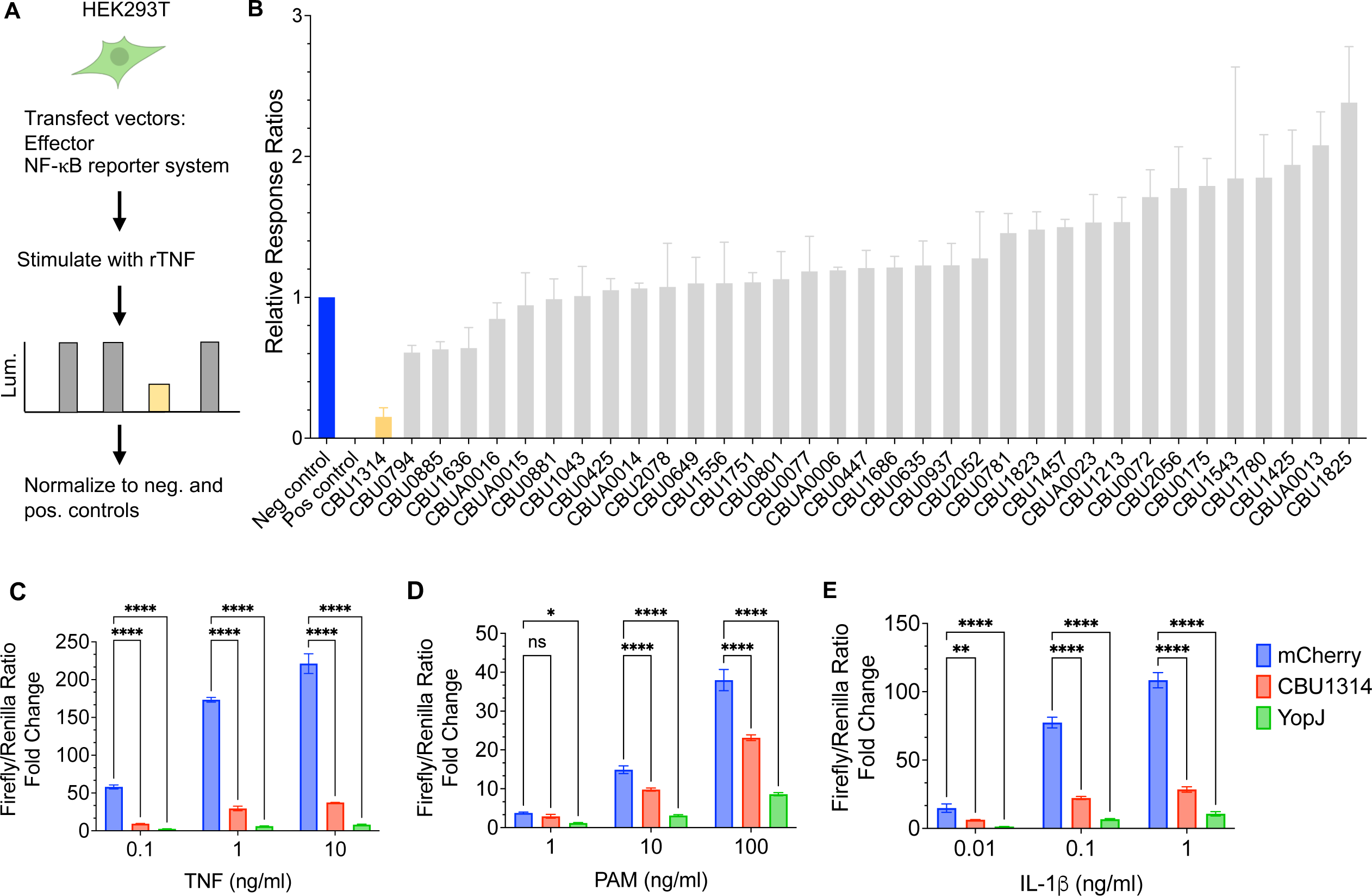
*C. burnetii* effector CBU1314 suppresses NF-κB signaling downstream of various agonists. HEK293T cells were transfected with each effector-encoding vector as indicated (pgLAP1) and the NF-κB luciferase reporter system. For Pam3CSK4 (PAM) experiment, cells were additionally transfected with a TLR2-encoding vector. A) Schematic diagram of screen. Graphical cell created with BioRender.com. B) Cells were stimulated with TNF at 1ng/ml for 4hrs. Data shown are combined from two screens, with triplicate replicates each. Cells were stimulated with TNF (C) or PAM (D) for 4hrs, or IL-1β (E) for 8hrs prior to the luciferase assay. C-E) Fold change was calculated by dividing the Firefly/Renilla luciferase ratio of TNF-stimulated cells by the Firefly/Renilla luciferase ratio of unstimulated cells. Bar graphs display the mean +/- SD of triplicate replicates. Data is representative of 3 independent experiments for C-E. Statistical analysis for C-E: Two-way ANOVA with Dunnett’s multiple comparisons test within each concentration was conducted.

CBU1314 was identified as a translocated effector using *Legionella pneumophila* as a surrogate host (Chen et al., 2010), and subsequently confirmed to be injected by *C. burnetii* into host cells (Weber et al., 2016). Importantly, CBU1314 suppressed NF-κB activation in a dose-dependent manner (Supplemental Fig. 1). Furthermore, CBU1314 potently suppressed NF-κB activation in response to increasing amounts of TNF, nearly to the same extent as YopJ, a *Yersinia* secreted effector protein that potently suppresses IKK activation (Fig. 2C) (Mukherjee et al., 2006; Orth et al., 2000; Zhou et al., 2005). In addition to TNF signaling, CBU1314 suppressed NF-κB activation downstream of other inflammatory activators, including the TLR2 agonist Pam3CSK4 or the cytokine IL-1β (Fig 2D-E, Supplemental Fig. 2). Together, these data demonstrate that CBU1314 suppresses NF-κB-induced gene expression downstream of multiple innate immune receptors.

### CBU1314 modulates immune gene expression in THP-1 cells

To test the potential contribution of CBU1314 to disruption of innate immune gene expression in a more physiologically relevant immune cell type, we turned to the monocytic THP-1 cell line.

We next tested whether CBU1314 suppressed gene expression downstream of TNF signaling in THP-1 cells using lentiviral vectors encoding CBU1314, which was able to suppress NF-κB activation in HEK293T cells as expected (Supplemental Fig. 3A-B), mCherry as a negative control, or YopJ as a known inhibitor of NF-κB signaling. mRNA analysis of cells expressing CBU1314 revealed that TNF-mediated induction of the innate immune genes *IRG1, TNF, IL6,* and *IL12B* are significantly suppressed by CBU1314 and YopJ relative to mCherry-expressing control cells (Fig. 3A). Interestingly however, our mRNA analysis also revealed a subset of genes that were not suppressed or were further induced following TNF treatment by CBU1314, including *IL1A*, *IL1B*, *IL10* and *CXCL8* (Fig. 3B). These data indicate that this effector modulates rather than exclusively inhibits host gene expression in response to TNF signaling.

**Figure 3.**
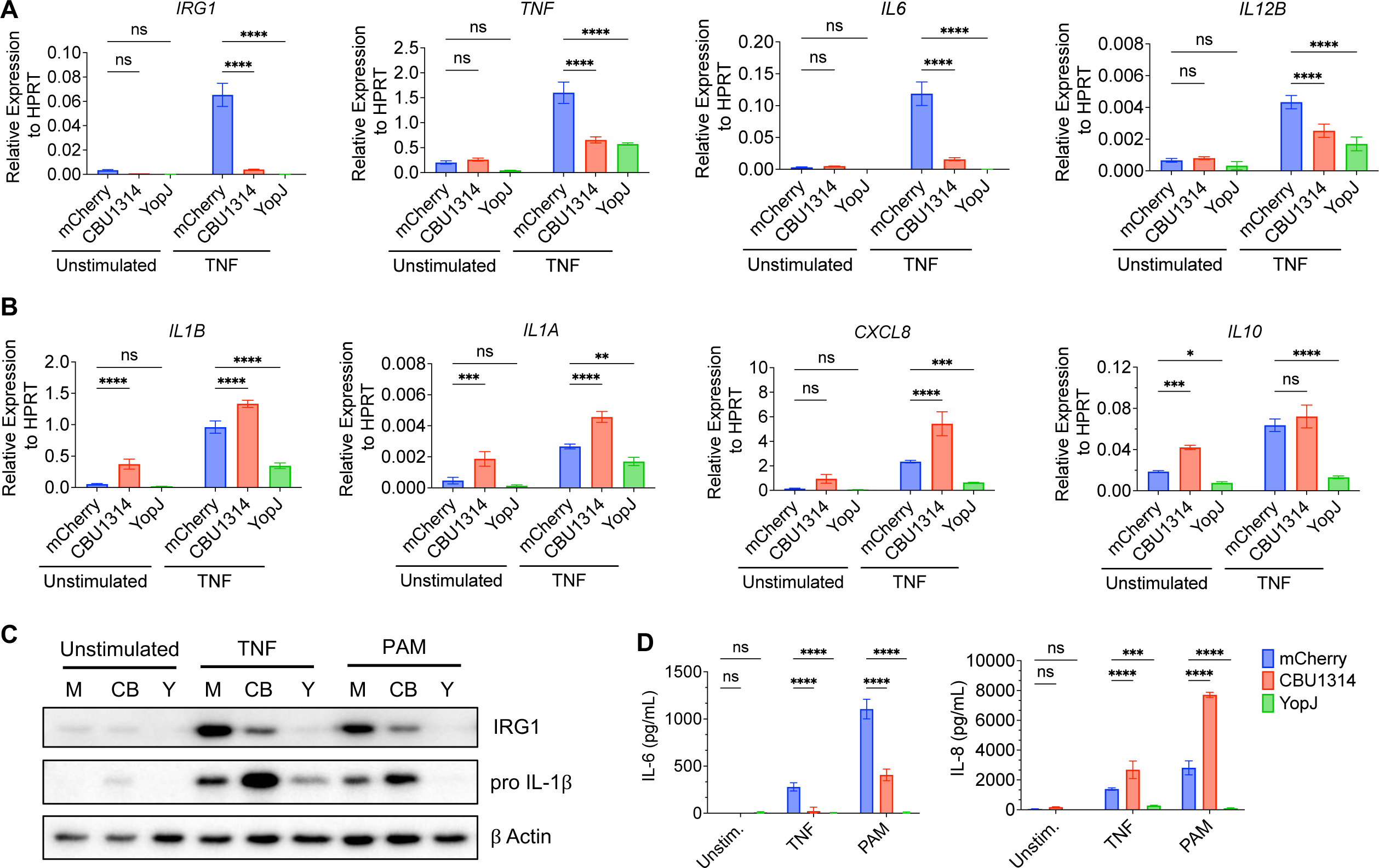
CBU1314 modulates the immune response in THP-1 cells. THP-1 cells expressing CBU1314 or controls (mCherry and YopJ) were left unstimulated or stimulated with TNF for 6hrs (A and B) or 24hrs (C and D). RNA was harvested at 6hrs for mRNA analysis (A and B), and protein and supernatants were harvested at 24hrs for Western blot (C) and ELISAs (D). Bar graphs display the mean +/- SD of triplicate replicates. All data is representative of 3 independent experiments. Statistical analysis for A and B: Two-way ANOVA with Dunnett’s multiple comparisons test for each unstimulated or TNF-stimulated group; for D: One-way ANOVA with Dunnett’s multiple comparisons test within each unstimulated or stimulated group.

Either suppression or induction of gene transcription would also be expected to affect their levels of protein production and, in the case of cytokines and chemokines, their subsequent secretion from cells. CBU1314-expressing THP-1 cells stimulated for 24 hours with TNF showed decreased IRG1 and increased pro-IL-1β protein levels relative to mCherry-expressing control cells, in agreement with the mRNA expression data (Fig. 3C). CBU1314 expression in THP-1 cells also led to decreased IL-6 and increased IL-8 secretion in response to TNF treatment relative to control cells, in agreement with decreased *IL6* mRNA levels and increased mRNA levels of *CXCL8,* which encodes IL-8 (Fig. 3D). Consistently, Pam3CSK4 treatment, which stimulates TLR2 signaling, led to increased pro-IL-1β protein levels and IL-8 secretion from CBU1314-expressing cells relative to control cells, and decreased IRG1 protein levels and IL-6 secretion, similar to TNF stimulation (Fig. 3C-D). This contrasted with THP-1 cells expressing the *Yersinia* effector YopJ, which exhibited broad suppression of IRG1, pro-IL-1β, IL- 6, and IL-8 production in response to TNF or Pam3CSK4 stimulation relative to mCherry-expressing control cells (Fig. 3C-D). Together, these results show that CBU1314 modulates the expression and secretion of a subset of innate immune proteins in THP-1 cells.

### CBU1314 suppresses both NF-κB and MAPK signaling in THP-1 cells

To determine whether CBU1314 suppresses activation of RelA or other components of NF-κB signaling, we ectopically expressed CBU1314 in THP-1 cells and performed Western blots for various markers of NF-κB pathway activation. In resting state, NF-κB is held sequestered in the cytosol by the inhibitory protein IκBα. Pathway activation leads to a phosphorylation cascade that activates the IKK kinase complex, which phosphorylates IκBα and targets it for proteasomal degradation. NF-κB is also phosphorylated by various kinases, leading to optimal NF-κB activation. Once IκBα is degraded, NF-κB can enter the nucleus and act as a transcription factor (Dorrington & Fraser, 2019; Mitchell, Vargas, & Hoffmann, 2016; Viatour, Merville, Bours, & Chariot, 2005). In contrast to THP-1 cells expressing the negative control mCherry, THP-1 cells expressing CBU1314 displayed substantially decreased phosphorylation of the kinases IKKα/β and the NF-κB subunit RelA downstream of TNF signaling. There was also a defect in IκBα degradation (Fig. 4A). YopJ-expressing cells also displayed decreased phosphorylation of proteins and degradation of IκBα, as expected for this *Yersinia* effector which disrupts cytosolic post-translational modifications that are required for signaling (Fig. 4A) (Mukherjee et al., 2006; Orth et al., 2000; Zhou et al., 2005). These results suggest that CBU1314 dampens early NF-κB pathway activation events that occur in the cytosol of host cells.

**Figure 4.**
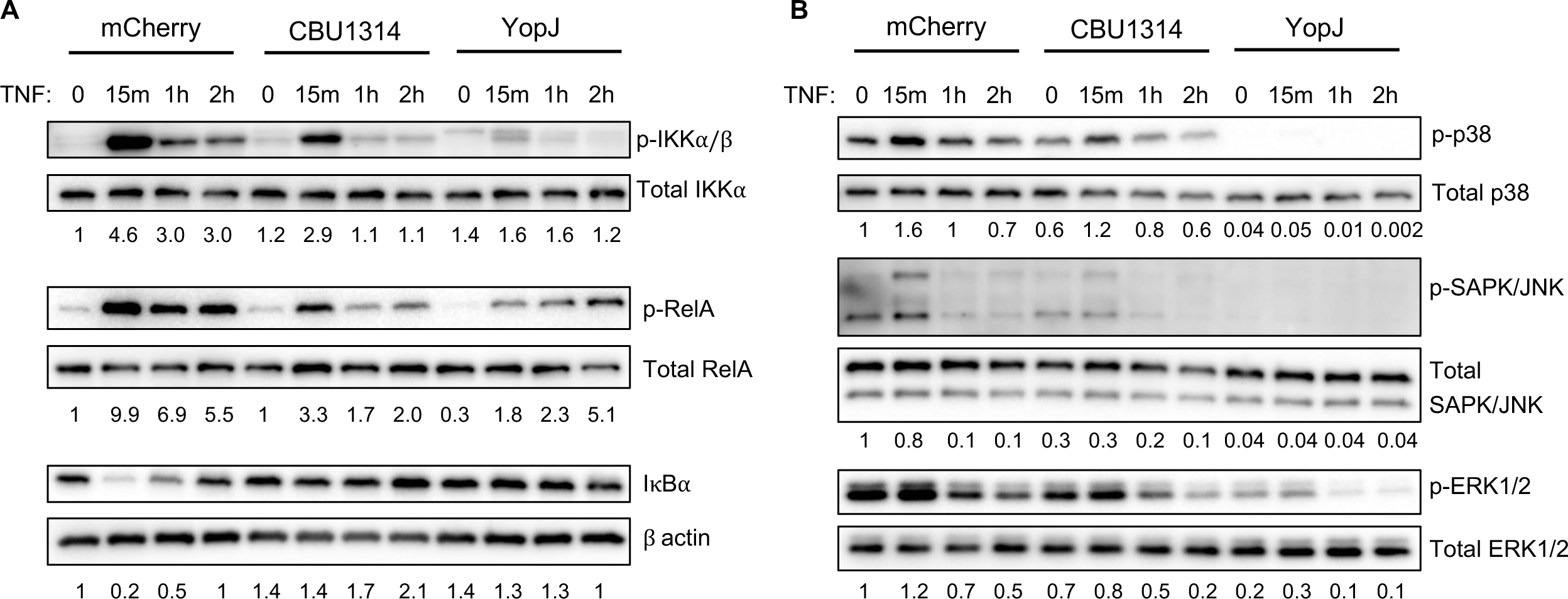
CBU1314 suppresses the activation of NF-κB and MAPK signaling components. THP-1 cells expressing mCherry, CBU1314, or YopJ were stimulated with TNF for the indicated times, then harvested for protein analysis for NF-κB (A) and MAPK (B) proteins. Numbers under blots indicate quantification ratios for each target relative to its control and normalized to mCherry 0 TNF lanes (set to 1). Data is representative of 3 independent experiments.

In addition to NF-κB, TNF also activates other signaling pathways, such as the MAPK family. MAPKs are activated downstream of several different receptors and similar to NF-κB, they promote many different responses, such as cell survival, growth and inflammation (W. Zhang & Liu, 2002). THP-1 cells expressing CBU1314 revealed slightly decreased phosphorylation of p38, ERK1/2, and JNK MAPKs at the 15-minute timepoints following TNF stimulation, relative to control mCherry-expressing cells, albeit in a more modest manner compared to the suppression we observed of the NF-κB pathway (Fig. 4B). In contrast, THP-1 cells expressing YopJ exhibited a substantial decrease in p38, ERK1/2, and JNK MAPK phosphorylation, as expected (Mukherjee et al., 2006; Orth et al., 2000; Zhou et al., 2005). Overall, these results suggest that CBU1314 dampens the activation of NF-κB and MAPK signaling downstream of TNF.

### CBU1314 localizes to the nucleus and associates with chromatin in THP-1 cells

Notably, CBU1314 dampening of cytosolic NF-κB and MAPK signaling events is relatively modest compared to YopJ suppression (Fig. 4), yet the ability of CBU1314 to suppress immune gene expression was similar to that of YopJ (Fig. 3A). This apparent disconnect between the effect of CBU1314 on signal transduction events and the ultimate outcome of gene expression raised the question of whether suppression of cytosolic events was indeed the major driver of gene downregulation.

CBU1314 localizes to the host cell nucleus, where it associates with chromatin and modulates host transcription (Chen et al., 2010; Weber et al., 2016). In agreement with previous findings (Weber et al., 2016), we found that GFP-tagged CBU1314 colocalized mostly with the nuclear fraction of HEK293T cells, with some cytoplasmic localization (Supplemental Fig. 4A). However, when ectopically expressed in THP-1 cells, we found that CBU1314 is exclusively localized in the nucleus and associates tightly with chromatin, appearing only in the chromatin-associated fractions extracted with the highest salt concentrations (Supplemental Fig. 4B). Altogether, these findings indicate that CBU1314 also localizes to the nucleus in innate immune cells and suggests that it exerts its function exclusively from that host compartment.

### CBU1314 downregulates mRNA expression of several TNFR1 signaling pathway proteins

As CBU1314 associates with host chromatin and a previous study indicated that CBU1314 modulates gene transcription (Weber et al., 2016), we hypothesized that CBU1314 might indirectly inhibit these cytosolic events by suppressing transcription of genes encoding immune signaling components themselves. We focused on TNFR1 signaling proteins upstream of our identified CBU1314-mediated block in IKK phosphorylation and IκBα degradation. Since HEK293T cells expressing CBU1314 also displayed a block in IκBα degradation (Supplemental Fig. 3C), we used these cells to test our hypothesis. Intriguingly, mRNA levels of critical components of the TNFR1 membrane-proximal signaling complex, including *TNFRSF1A* (encoding TNFR1 itself), *TRADD*, *TRAF2, RIPK1, SHARPIN, RNF31* (encoding HOIP)*, TAB1, IKBKG* (encoding NEMO), and *BIRC2* and *BIRC3* (encoding cIAP1 and cIAP2, respectively) were all significantly downregulated by CBU1314 in TNF-stimulated cells compared to negative control cells, while *MAP3K7* (encoding TAK1) was unaffected (Supplemental Fig. 5). Notably, expression of the majority of these genes was either unaffected or affected to a lesser extent in cells expressing the *Yersinia* effector YopJ compared to CBU1314, indicating that suppression of immune signaling by these two bacterial effectors is mechanistically distinct. The downregulation of many of these genes by CBU1314 was also seen in unstimulated cells.

**Figure 5.**
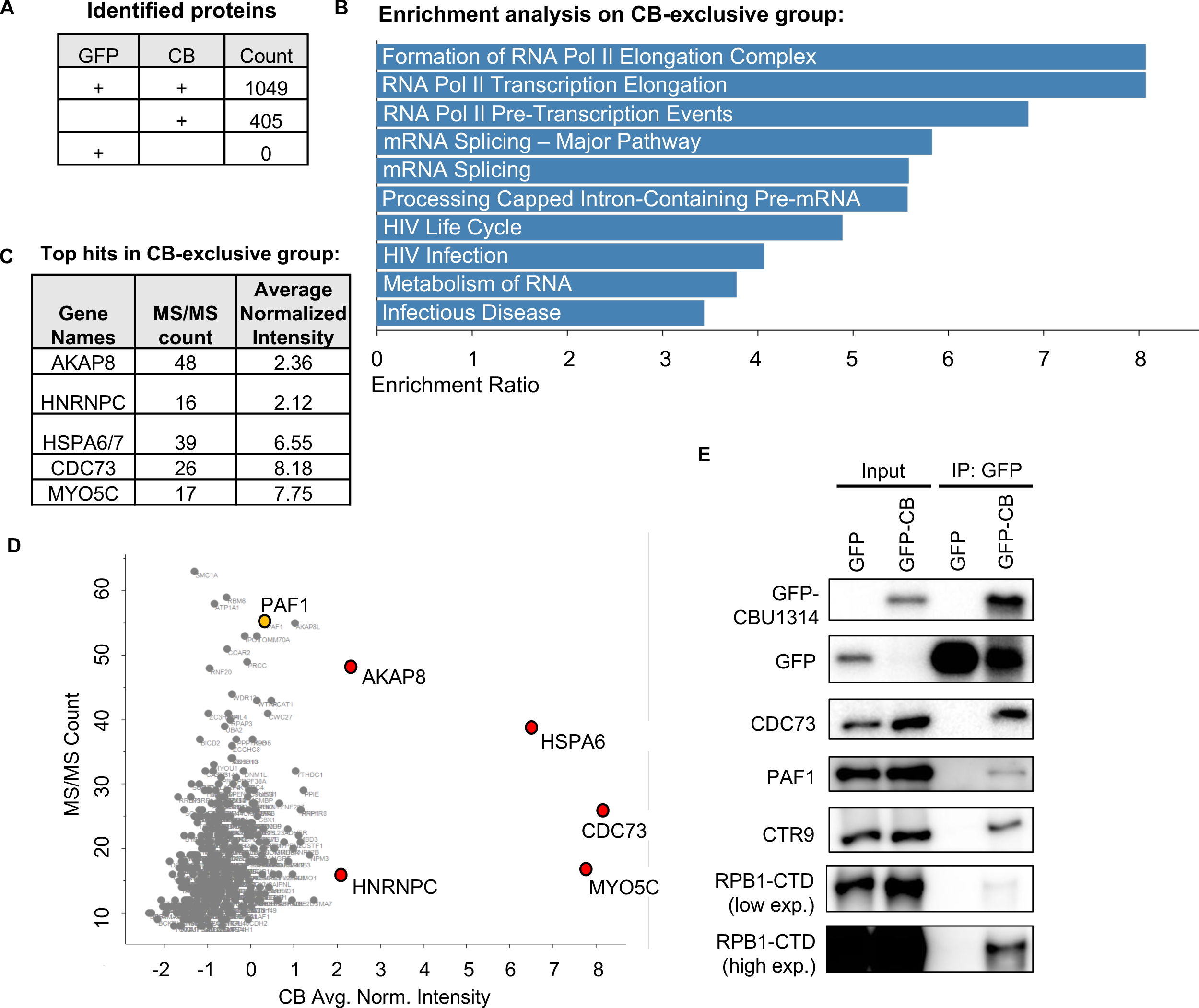
IP-MS and co-IP experiments reveal interaction of CBU1314 with the host PAF1 complex. A) Total number of identified proteins in GFP or CBU1314 (CB) IP samples. B) Reactome pathway enrichment analysis conducted on the set of proteins that were only identified in the CBU1314 samples (CB-exclusive group). C) Table of top candidates from the CB-exclusive group. D) Plot of MS/MS count vs. average normalized intensity of CB-exclusive proteins, with top hits highlighted in red, and PAF1 highlighted in yellow. E) Co-immunoprecipitation in HEK293T cells expressing GFP alone or GFP-CBU1314. Members of the PAF1 complex and RPB1-CTD were detected via immunoblot analysis.

Altogether, our results indicate that CBU1314 broadly inhibits the expression of TNFR1 and NF- κB signaling components, which may contribute to suppression of cytosolic signaling events.

### Immunoprecipitation-mass spectrometry identifies PAF1 complex components as host binding partners of CBU1314

We next investigated whether CBU1314 interacts with any host protein partners that could give a clue to its mechanism of action. Yeast genetic interaction profiling previously predicted potential host genes, complexes, and/or pathways modulated by bacterial effectors, including CBU1314 (Patrick et al., 2018). Intriguingly, CBU1314 was predicted from this analysis to interact with nuclear activities and functions, including the yeast mediator of RNA polymerase II transcription (Mediator) complex, chromatin remodeling, and pre-mRNA splicing proteins (Patrick et al., 2018). However, the specific host targets of CBU1314 in mammalian cells remain unknown. To specifically identify such targets, we performed immunoprecipitation (IP)-mass spectrometry (MS) to find potential host protein partners of CBU1314. MS analysis revealed a large subset of host proteins that were identified in both control and experimental sample groups, and a subset that were only present in the GFP-CBU1314 samples (Fig. 5A). Pathway enrichment analysis of the differentially identified proteins between the two sample groups revealed enrichment of mRNA splicing and processing functions, among other RNA-related functions (Supplemental Fig. 6). Further pathway enrichment analysis of the proteins identified only in the CBU1314 group also revealed enrichment of RNA- and transcription-associated functions, including formation of the RNA Pol II elongation complex, transcription elongation, and mRNA splicing (Fig. 5B). These results are in line with the previously described function of CBU1314 in modulating host transcription (Weber et al., 2016) and the predicted pathway interactions in the yeast genetic system (Patrick et al., 2018).

**Figure 6.**
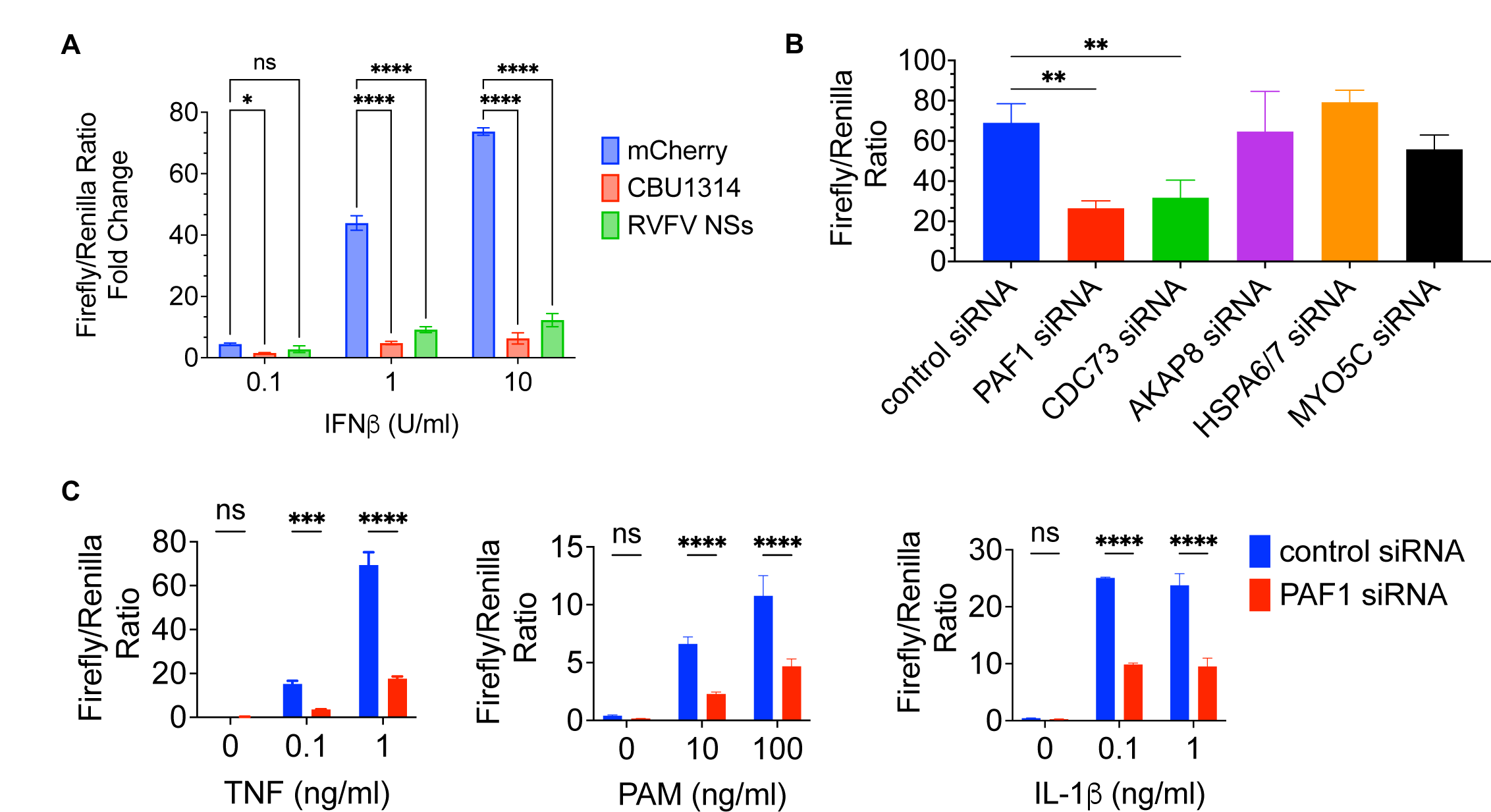
CBU1314 suppresses type I IFN signaling, and PAF1C promotes NF-κB- dependent transcription. A) CBU1314 or controls mCherry and Rift Valley Fever Virus NSs (RVFV NSs) were expressed in HEK293T cells, along with an ISRE-luciferase reporter system. Cells were stimulated with the indicated concentrations of IFNβ for 8hrs prior to the luciferase assay. Results are displayed as fold change over luciferase ratio readings for unstimulated cells. B) siRNA knockdown of the indicated targets was combined with the NF-κB luciferase assay in HEK293T. Cells were stimulated with TNF at 1ng/ml for 4hrs prior to measuring luciferase activity. C) *PAF1* siRNA knockdown was combined with the NF-κB luciferase assay as in B. Cells were stimulated with the indicated agonists prior to the luciferase assay. Graphs display the mean +/- SD of triplicate replicates. Data is representative of 2 independent experiments for A and B, and 3 for C. Statistical analysis for A: Two-way ANOVA with Dunnett’s multiple comparisons test within each concentration; for B: One-way ANOVA with Tukey’s multiple comparison’s test, with only significant comparisons shown; for C: Two-way ANOVA with Sidak’s multiple comparisons test at each agonist concentration.

The top five hits present in the CBU1314-exclusive group were AKAP8, HNRNPC, HSPA6/HSPA7, CDC73, and MYO5C (Fig. 5C). These proteins had a high average normalized intensity and/or MS/MS counts across all 4 IP replicates (Fig. 5D). Intriguingly, CDC73 is a member of the nuclear polymerase-associated factor 1 (PAF1) complex (PAF1C), which contains several proteins (PAF1, CDC73, CRT9, SKI8, LEO1). PAF1, LEO1, and SKI8 were also identified as potential binding partners of CBU1314 by IP-MS. PAF1C is involved in various transcriptional functions, including transcription elongation by RNA Pol II (Francette, Tripplehorn, & Arndt, 2021; Hou et al., 2019; Kim et al., 2010; Yu et al., 2015). PAF1C also promotes the expression of immune genes (Kenaston et al., 2022; Marazzi et al., 2012; Parnas et al., 2015; Petit et al., 2021). Furthermore, both the influenza virus NS1 and dengue virus NS5 proteins interact with and antagonize PAF1C, leading to suppression of antiviral type I interferon-stimulated gene (ISG) expression (Marazzi et al., 2012; Petit et al., 2021; Shah et al., 2018). Altogether, these findings suggested that CBU1314 might be a bacterial modulator of PAF1C function in order to suppress immune signaling.

To directly test whether CBU1314 interacts with components of the PAF1C, we performed co-IP experiments. Consistent with the IP-MS findings, CBU1314 pulled down the PAF1C members CDC73, PAF1, and CTR9, as well as the RNA Polymerase II subunit RPB1-CTD (Fig. 5E) in TNF-stimulated cells. However, these interactions were not dependent on TNF signaling, as CBU1314 also pulled down CDC73 and RPB1-CTD in unstimulated cells (Supplemental Fig. 7A). Importantly, co-IP studies from purified nuclear fractions demonstrated that only CBU1314, and not GFP alone, interacted with CDC73 in the nucleus, highlighting the specificity of the interaction between CBU1314 and components of the PAF1 complex (Supplemental Fig. 7B).

**Figure 7.**
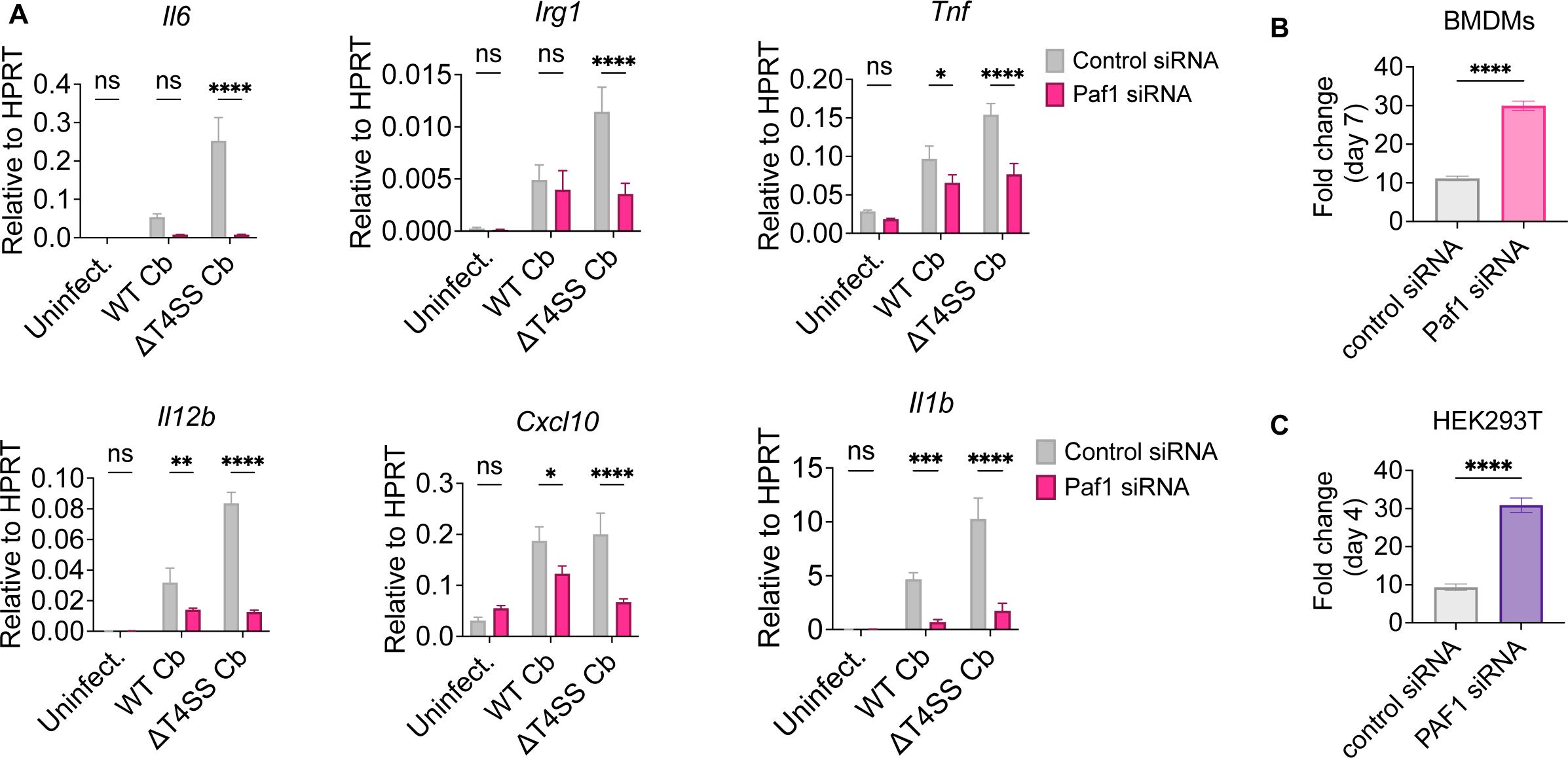
PAF1 promotes gene expression during infection and restricts intracellular bacterial replication. A) Murine bone marrow-derived macrophages (BMDMs) were treated with *Paf1* siRNA for 48hrs, then infected with WT or T4SS-mutant *C. burnetii* (ΔT4SS Cb) at an MOI of 50 for 24hrs. Cell lysates were harvested for mRNA expression analysis. B-C) siRNA knockdown of (B) *Paf1* in BMDMs or (C) *PAF1* in HEK293T cells. Cells were infected with mCherry-expressing WT *C. burnetii* at MOI 100. Samples were harvested at day 0 and 7 for (B), and day 0 and 4 for (C), and fold change was calculated based on genome equivalents. Bar graphs display the mean +/- SD of triplicate replicates. Data is representative of 2-3 independent experiments. Statistical analysis for A: Two-way ANOVA with Sidak’s multiple comparisons test at each agonist concentration or infection condition; for B and C: Unpaired t- test.

### CBU1314 inhibits type I interferon signaling, and PAF1C also promotes NF-κB-dependent transcription

Since CBU1314 associates with PAF1C, and this complex is important for IFN-dependent gene expression (Marazzi et al., 2012), we asked whether CBU1314 could also suppress IFN-dependent gene expression. We found that CBU1314 potently inhibited ISRE-dependent luciferase induction downstream of IFNβ relative to mCherry-expressing control cells, to an extent similar to the Rift Valley fever virus (RVFV) nonstructural (NS) protein encoded on the S-segment (NSs), which is a known potent inhibitor of host transcription (Fig. 6A) (Wuerth & Weber, 2016).

Next, we further investigated the role of PAF1C in immune signaling. In addition to its role in IFN-dependent gene expression, PAF1C contributes to TNF expression downstream of TLR4 in murine DCs (Parnas et al., 2015), and the complex member CTR9 promotes IL-6-dependent gene expression (Youn et al., 2007). A recent study indicated that the promoters of PAF1- dependent genes downstream of a variety of different immune stimuli had motifs for several transcription factors, including the NF-κBs c-Rel and RelA, and the MAPK-regulated factor c-Jun (Kenaston et al., 2022). These findings indicate that PAF1C broadly regulates immune gene expression downstream of receptors that activate these transcription factors. We thus examined whether knockdown of PAF1 could affect NF-κB-mediated gene transcription. siRNA knockdown of the top MS candidates combined with the TNF-induced NF-κB luciferase assay in HEK293T cells revealed that only knockdown of the PAF1C complex members *PAF1* and *CDC73* led to a significant decrease in NF-κB-dependent luciferase signal compared to control siRNA-treated cells, whereas knockdown of *AKAP8*, *HSPA6/7*, or *MYO5C* did not (Fig. 6B).

Furthermore, *PAF1* knockdown followed by stimulation with a range of TNF concentrations and other agonists revealed that PAF1 was required for maximal NF-κB-dependent luciferase induction in response to TNF, the TLR2 agonist Pam3CSK4, and IL-1β (Fig. 6C). These results suggest that the PAF1 complex not only participates in IFN-dependent gene expression, but also mediates NF-κB-dependent gene expression downstream of several innate immune receptors, in agreement with the predictions by a recent study (Kenaston et al., 2022).

Collectively, our results indicate that CBU1314 inhibits gene expression downstream of multiple innate immune pathways, and that the host factor PAF1C promotes those very same pathways, raising the possibility that CBU1314 may suppress immune gene expression by antagonizing PAF1C, similarly to the influenza NS1 and dengue NS5 viral proteins (Marazzi et al., 2012; Petit et al., 2021).

### PAF1 contributes to gene expression downstream of cytokine signaling and during *C. burnetii* infection

We next asked whether PAF1 controls immune gene expression in THP-1 monocytes in response to cytokine stimulation or during *C. burnetii* infection. A previous study in THP-1 cells revealed that PAF1C promotes RNA polymerase II pause release and correlates with increased mRNA levels at various genes, suggesting a role in gene expression (Yu et al., 2015), but this study was performed in the absence of immune stimulation. We found that siRNA-mediated silencing of *PAF1* in THP-1 cells followed by stimulation with either TNF or IFNβ resulted in modest decreased expression of the TNF-stimulated genes *CXCL8* and *IL1B* and the ISGs *RSAD2* (encoding viperin) and *OAS1* compared to control siRNA-treated cells (Supplemental Fig. 8A-B). We next infected control and *PAF1* siRNA-treated THP-1 cells with WT *C. burnetii*.

*C. burnetii* infection led to a modest increase in immune gene expression compared to uninfected cells, and *PAF1* knockdown slightly decreased expression of some genes (Supplemental Fig. 8C). In contrast, in THP-1 cells infected with the T4SS-mutant *C. burnetii,* which are unable to translocate effectors and thus cannot suppress immune gene expression, we observed a large increase in gene expression compared to uninfected or WT *C. burnetii-* infected cells. Furthermore, *PAF1* knockdown led to a substantial decrease in gene expression of several genes (Supplemental Fig. 8C). We also found that *Paf1* contributed to *C. burnetii*- induced immune gene expression in primary murine bone marrow-derived macrophages (BMDMs), where *Il6*, *Irg1*, *Tnf*, *Il12b*, *Cxcl10* and *Il1b* showed decreased expression in *Paf1* siRNA- compared to control siRNA-treated cells (Fig. 7A). These data indicate that PAF1 is required for maximal immune gene expression in response to TNF and type I IFN and during *C. burnetii* infection of human monocytes and murine macrophages.

### PAF1 contributes to restriction of intracellular *C. burnetii* replication

PAF1C restricts replication of several viruses, including HIV, influenza, and dengue virus (Abdel-Mohsen et al., 2015; Liu et al., 2011; Marazzi et al., 2012; Petit et al., 2021; Raposo et al., 2013; Shah et al., 2018), yet its potential role in bacterial restriction has never been examined. We therefore investigated whether PAF1 plays a role in restricting intracellular *C. burnetii* replication. We first examined murine BMDMs because they are highly restrictive of *C. burnetii* replication in a TLR2- and TNF-dependent manner (Bradley et al., 2016; Zamboni et al., 2004), and thus could provide a robust system to measure any increase in permissiveness of bacterial replication. Knockdown of *Paf1* in BMDMs led to a significant increase in bacterial replication, from 10-fold growth in control siRNA-treated cells to almost 30-fold growth in *Paf1* siRNA-treated cells at day 7 post-infection (Fig. 7B). Similarly, HEK293T cells exhibited significantly increased bacterial replication in *PAF1* siRNA-treated cells compared to control siRNA-treated cells at day 4 post-infection (Fig. 7C). Overall, these results indicate that PAF1C is required to restrict intracellular *C. burnetii* replication.

## Discussion

We describe here the discovery of *a C. burnetii* effector, CBU1314, that suppresses innate immune signaling and gene expression. Our data indicate that CBU1314 inhibited NF-κB and MAPK signaling and gene expression downstream of several immune receptors, including TNF, TLR2, and IL-1R, and also inhibited type I IFN-dependent gene expression. These results indicate that CBU1314 is a broadly-acting suppressor of multiple innate immune pathways.

Expression of CBU1314 in THP-1 cells led to the modulation of innate immune genes downstream of TNF, where some genes were suppressed and others were induced. Interestingly, immune responsive gene 1 (*IRG1*), which was suppressed by CBU1314, is a potent inhibitor of bacteria through production of itaconic acid (Luan & Medzhitov, 2016; Michelucci et al., 2013; Naujoks et al., 2016), and a previous study identified a ChIP-seq binding peak for CBU1314 close to this gene (Weber et al., 2016). Whether the binding of CBU1314 to the *IRG1* locus leads to inhibition of *IRG1* expression warrants further study.

Given that CBU1314 potently inhibited NF-κB-dependent luciferase induction, we were surprised to find NF-κB-regulated genes that were not inhibited by this effector. However, these genes are not solely regulated by NF-κB (DeLaney et al., 2019; Palazon, Goldrath, Nizet, & Johnson, 2014), and it is possible that other transcription factors not inhibited by this effector, or activated by CBU1314 activity through effector-triggered immunity (Lopes Fischer, Naseer, Shin, & Brodsky, 2020), contribute to gene expression. Furthermore, we found that CBU1314 associates with PAF1C, and PAF1C was recently described as promoting the expression of some immune genes and suppressing others (Kenaston et al., 2022). Thus, it is possible that the disruption of PAF1C function by CBU1314 inversely modulates the induction or suppression of immune genes regulated by PAF1C. Future studies to investigate the global effects of CBU1314 on gene expression and experiments to elucidate the role of PAF1C or other host factors will provide insight into this differential gene expression.

Our studies revealed that CBU1314 led to suppressed phosphorylation of IKK and the NF-κB subunit RelA, and reduced degradation of the inhibitory protein IκBα. We also found that CBU1314 led to dampened phosphorylation of MAPKs, although this phenotype was more subtle compared to what we observed for NF-κB signaling. These results were unexpected because these signaling events occur in the cytosol, whereas both our findings and those of others indicate that CBU1314 localizes to the nucleus (Chen et al., 2010; Weber et al., 2016). We therefore hypothesized that CBU1314 indirectly suppresses these signaling events as a consequence of its effect on transcription, potentially of key components of immune signal transduction pathways. Indeed, the expression of multiple TNFR1 pathway components was downregulated by CBU1314. These results provide a possible mechanism for how a nuclear protein that modulates host transcription could suppress cytosolic signaling events: by decreasing the expression of upstream regulators that are important for activating this pathway.

Our IP-MS and subsequent experiments revealed that CBU1314 interacts with members of PAF1C, which is linked to the regulation of Pol II-mediated transcription elongation, mRNA processing and alternative cleavage/polyadenylation, chromatin modification, expression of antiviral genes, and restriction of viral replication (Abdel-Mohsen et al., 2015; Francette et al., 2021; Hou et al., 2019; Kim et al., 2010; Liu et al., 2011; Marazzi et al., 2012; Raposo et al., 2013; Yang et al., 2016; Yu et al., 2015). Interestingly, a study that used yeast genetics to predict interactors with bacterial effectors identified a possible genetic interaction between PAF1C and a different *C. burnetii* effector, CBU0794 (Patrick et al., 2018). CBU0794 was part of our initial screen and showed a slightly decreased Firefly/Renilla luciferase ratio compared to the negative control, but not to the same extent observed for CBU1314. Future studies testing this effector’s predicted interaction with PAF1C and investigating whether it collaborates with or antagonizes CBU1314 function would be of interest.

NS proteins from dengue and influenza viruses have been found to suppress innate immune signaling by inhibiting PAF1C (Marazzi et al., 2012; Petit et al., 2021; Shah et al., 2018), providing a shared strategy used by these viruses to evade host responses. Critically, we find that *PAF1* knockdown not only led to decreased expression of ISGs induced by *C. burnetii* infection, similar to what was previously described for viral infection (Marazzi et al., 2012), but also led to decreased NF-κB-dependent gene induction downstream of various receptors.

Because of CBU1314’s ability to suppress innate immune signaling and gene expression, and the similar effect on immune gene expression observed in cells following PAF1 knockdown or CBU1314 overexpression, we hypothesize that CBU1314 antagonizes PAF1C function. Future studies will be needed to further elucidate the interaction between CBU1314 and PAF1C and its possible role in modulating host transcription and the immune response. Intriguingly, it was recently shown that the *C. burnetii* effector AnkG interacts with the host factors DDX21 and 7SK snRNP complex, which are involved in host transcription (Cordsmeier et al., 2022). Thus, *C. burnetii* may employ multiple effectors that interact with host transcriptional machinery to modulate gene expression.

PAF1C has been previously linked to regulating ISG expression during viral infections (Marazzi et al., 2012; Petit et al., 2021; Shah et al., 2018), LPS-induced TNF expression (Parnas et al., 2015), IL-6-dependent gene expression (Youn et al., 2007), and more recently, gene expression in response to a variety of PRRs and cytokine stimuli (Kenaston et al., 2022). Our study shows a contribution for PAF1 specifically to NF-κB signaling and gene expression during bacterial infection and in response to various innate immune pathways, including TNFR, TLR2, and IL- 1R, in monocytes and macrophages. Furthermore, PAF1C is required to restrict the intracellular replication of several viruses, including HIV, influenza and dengue (Abdel-Mohsen et al., 2015; Liu et al., 2011; Marazzi et al., 2012; Petit et al., 2021; Raposo et al., 2013; Shah et al., 2018). Our data show that PAF1C is also required to restrict intracellular *C. burnetii* replication.

Collectively, these findings indicate that PAF1C is a central player in the regulation of IFN- and NF-κB-dependent immune responses and cell-intrinsic defense against a variety of intracellular pathogens.

Altogether, our studies reveal a nuclear *C. burnetii* effector, CBU1314, that interacts with the host PAF1 complex and broadly inhibits host signaling pathways and immune gene expression. Furthermore, our findings indicate that PAF1C regulates gene expression downstream of multiple innate immune pathways and during infection, and that PAF1C plays a role in restricting intracellular *C. burnetii* replication, highlighting for the first time the importance of this host complex in cell-intrinsic defense against a bacterial pathogen. Thus, our findings provide new insight into mechanisms by which *C. burnetii* suppresses anti-bacterial immune responses and the role of the host complex PAF1C in regulating anti-bacterial immunity.

## Methods

### Ethics statement

We obtained deidentified human monocytes from the Human Immunology Core at the University of Pennsylvania. This facility has IRB approval to collect peripheral blood mononuclear cells from anonymous healthy donors for research purposes and does not release identifying information on the donors. Thus, all research using these cells was performed in compliance with the requirements of the US Department of Health and Human Services and the Declaration of Helsinki.

### Cell culture

HEK293T cells were cultured in Dulbecco’s Modified Eagle’s Medium (DMEM, Corning) supplemented with 10% Fetal Bovine Serum (FBS), penicillin/streptomycin (P/S) (Corning), and L-glutamine (Corning). For luciferase assays, HEK293T cells were plated in DMEM without phenol red (Gibco) supplemented with 10% FBS, sodium pyruvate (Invitrogen), and L-glutamine. THP-1 cells were cultured in Roswell Park Memorial Institute Media 1640 (RPMI, Corning) supplemented with 10% FBS, 0.05mM 2-mercaptoethanol (Bio-Rad), and P/S. Human primary monocytes were differentiated with 50ug/ml Human Macrophage Colony Stimulating Factor (hM-CSF; Gemini Bio-Products #300-161P) for six days, in RPMI supplemented with 10% FBS, L-glutamine, and penicillin/streptomycin. At the time of experiments, cells were switched to media without antibiotics (replating media). Murine bone marrow-derived macrophages (BMDMs) were generated by culturing C57BL/6 bone marrow in DMEM supplemented with 20% FBS, GlutaMAX (Invitrogen), sodium pyruvate (Invitrogen), P/S, and 20ng/ml of recombinant murine M-CSF (Peprotech, #315-02) for 6-7 days prior to replating. At the time of replating, cells were lifted using PBS with 2mM EDTA, and resuspended in complete media with only 10% FBS and no antibiotics. All cells were grown at 37°C with 5% CO2.

Where indicated, cells were stimulated with the following recombinant proteins: human TNF (PeproTech, #300-01A-10UG); human IFNβ (PeproTech, #300-02BC); human IFNγ (R&D Systems, #285-IF-100); Pam3CSK4 (Invivogen, #tlrl-pms); human IL-1β (PeproTech, #200-01B- 2ug); murine TNF (PeproTech, #315-01A); IFNα (R&D Systems, #11200-1).

### C. burnetii infection

All *C. burnetii* strains used in this study were derived from Nine Mile phase II, strain RSA439, clone 4. Infections were carried out with WT *C. burnetii*, mCherry-expressing *C. burnetii* (for growth curves) (Beare et al., 2009), or an *icmL*::Tn mutant (Carey, Newton, Lührmann, & Roy, 2011) (referred in the manuscript as ΔT4SS) strains. The *C. burnetii* strains were grown for 7 days in ACCM-2 at 37°C in 2.5% O2 and 5% CO2 in a tri-gas incubator (New Brunswick Scientific). The bacterial genome equivalents were calculated as described below. Infections were performed using either freshly grown bacteria or from frozen bacterial stocks at an MOI of 20, 50, or 100 as indicated. For mRNA expression analysis, THP-1 monocytes were seeded at 4x10^5^ cells/well in a 48-well plate in RPMI and infected with *C. burnetii*. 24hrs later, cell lysates were harvested for mRNA analysis, as described below. For ELISA experiments, THP-1 monocytes were seeded at 2X10^5^ cells/well in a 48-well plate in replating media (complete media without antibiotics) and differentiated into macrophages for 24hrs with PMA at 200nM. The following day, the media was replaced with fresh replating media to remove the PMA and cells were infected with *C. burnetii* strains for 24hrs prior to harvesting the supernatants for ELISA analysis. For human monocyte-derived macrophage (hMDM) experiments, cells were harvested with trypsin and seeded on day 6 of differentiation at 1X10^5^ cells/well in a 48-well plate with 25ug/ml of hM-CSF. A few hours later, hMDMs were left untreated or primed overnight with rIFNγ (R&D Systems #285-IF-100) at 100U/ml. The following day, cells were infected and harvested as indicated above for THP-1 macrophages.

### Genome equivalence (GE) measurements for infection and growth curves

*C. burnetii* DNA was extracted from the axenic cultures using the GenElute Bacterial Genomic DNA Kit (Sigma-Aldrich, #NA2120). Genome equivalents (GE) were measured by qPCR, using a standard curve generated by a pGEM-T vector encoding the *C. burnetii dotA* gene. qPCR was carried out in a Bio-Rad CFX96 machine using SsoFast EvaGreen SYBR Mix (Bio-Rad, #1725212) and *dotA* primers are listed in Table 1.

**Table 1.**
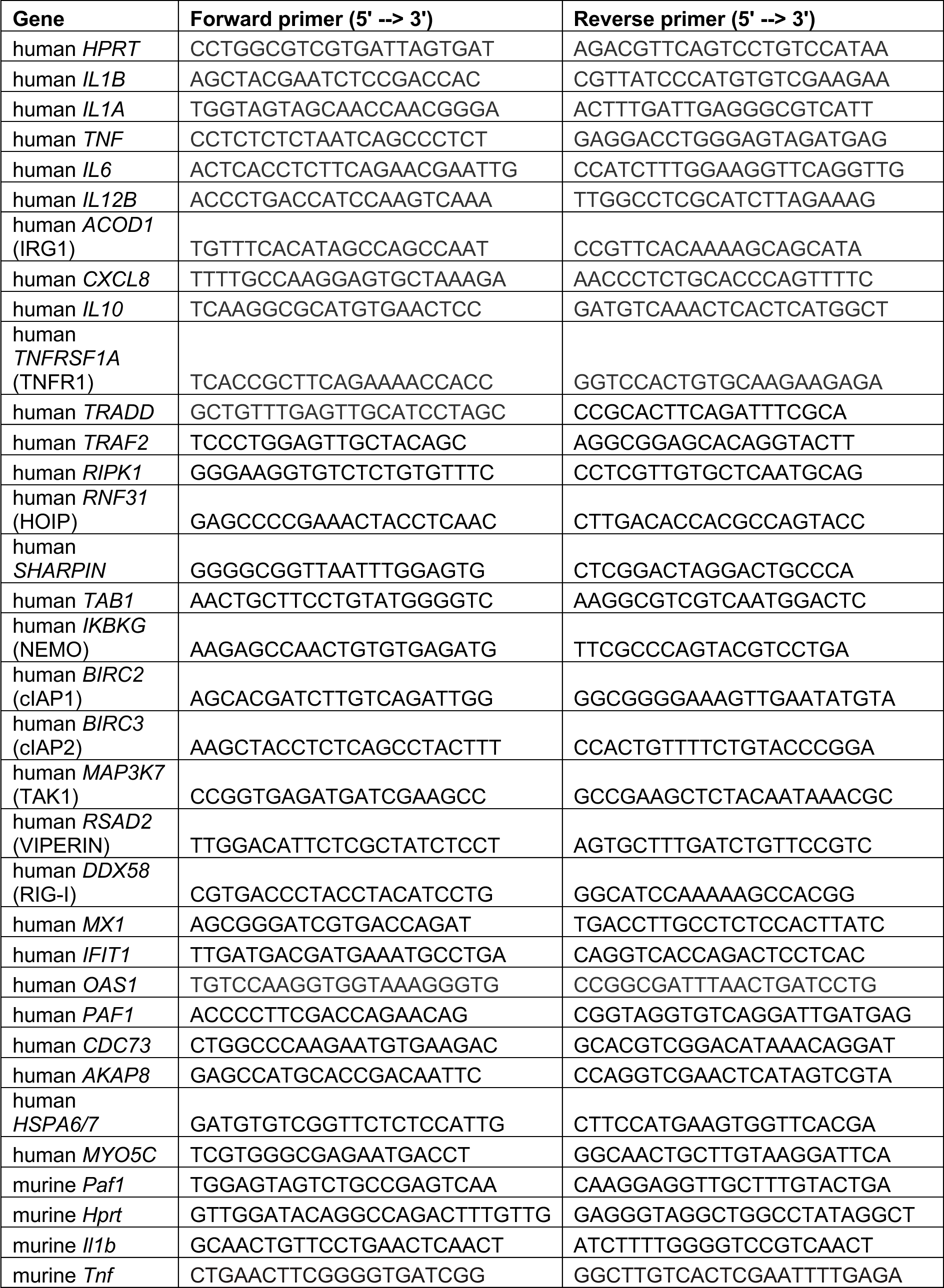
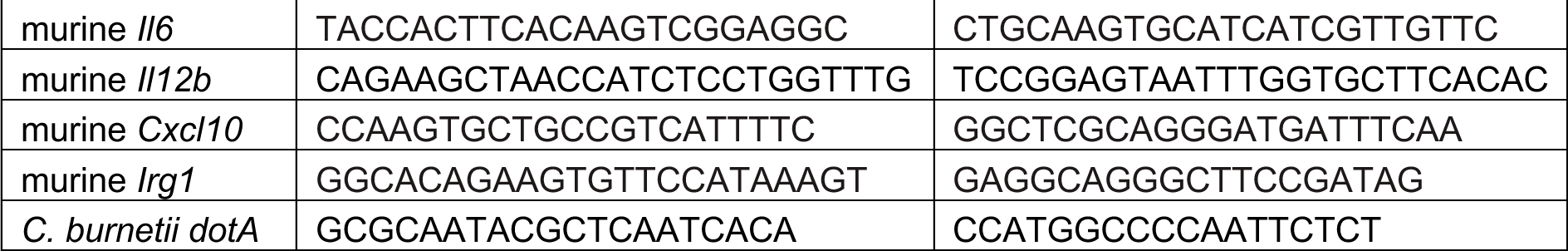
List of primers used in this study

### Vectors

Each effector or control gene was cloned into pENTR3C (Invitrogen #A10464) and transferred to pgLAP1 (Addgene #19702; deposited by Peter Jackson) using the Gateway system, in order to generate GFP N-terminally tagged proteins. For ectopic expression of genes in THP-1 cells, genes were synthesized into pTwist Lenti SFFV Puro WPRE vector with a C-terminal Streptag II (Twist Bioscience) for constitutive expression. The VSV-G and psPAX2 packaging plasmids (obtained from Paul Bates, University of Pennsylvania) were used to produce lentiviral particles. For the luciferase assays, the following plasmids were used: chicken β-actin promoter-driven *Renilla* luciferase (obtained from Elizabeth Kopp, Yale University), pGL4.32 NF-κB-Luc (Promega), pGL4.45 ISRE-Luc (Promega), and hTLR2 Flag (Addgene #13082; deposited by Ruslan Medzhitov). As a negative control, mCherry-encoding pgLAP1 was used, and as a positive control for NF-kB luciferase assay, YopJ-encoding pgLAP1 was used. As a positive control for the ISRE luciferase assay, a pcDNA3.1 vector encoding the Rift Valley Fever Virus NSs protein tagged with a Strep-tag was used (obtained from Kellie Jurado, University of Pennsylvania).

### Ectopic expression of effectors in HEK293T cells for mRNA analysis

Unless otherwise stated, HEK293T cells were seeded at 2x10^5^ cells/well of 48-well plate and transfected with 300ng of plasmid per well using Lipofectamine 2000 transfection reagent (Invitrogen, #11668019), as directed by the manufacturer. The next day, cells were left unstimulated or stimulated with 10ng/ml of TNF for 6hrs prior to harvest of cell lysates for mRNA analysis of TNFR1 pathway genes.

### Ectopic expression of effectors in THP-1 cells

Lentiviral particles were created in HEK293T cells as follows: cells were seeded in 10cm plates at 4x10^6^ cells/plate in HEK293T media without antibiotics. 24 hours later, cells were transfected with 1ug of VSV-G and 4ug of psPAX2 packaging plasmids, and 5ug of expression plasmid (pTwist Lenti), using Lipofectamine 2000 transfection reagent as directed by the manufacturer. Early the next morning, the media was replaced to remove the Lipofectamine. Cells were incubated at 37°C and 5% CO2 for 48hrs to allow for production of viral particles. THP-1 cells were transduced as follows: THP-1 cells were seeded at a concentration of 1x10^6^ cells/ml, 2ml per well of a 6-well plate in THP-1 media without antibiotics. Viral media from HEK293T cells was filtered through 0.45um filter and added to the THP-1 cells, 2ml per well. Polybrene transfection reagent (EMD Millipore, TR-1003-G) was added to each well at a final concentration of 8ug/ml. Cells were spun at 1250xg for 90 minutes at 30°C, then incubated for 1-2 days at 37°C with 5% CO2 prior to downstream experiments. Supernatants were harvested for ELISAs at 24hrs post-stimulation, cell lysates for mRNA expression were harvested at 6hrs post stimulation, and cell lysates for protein analysis were harvested as indicated in each figure.

### qPCR for RNA expression

RNA was extracted using the RNeasy Mini Plus Kit (Qiagen #74136), as directed by the manufacturer. cDNA was created using SuperScript II Reverse Transcriptase (Invitrogen #18- 064-071), as directed by the manufacturer, and by incubating at 42°C for 1hr. qPCR was conducted using SsoFast EvaGreen Supermix with Low ROX (Bio-Rad #1725212) using a Bio-Rad CFX96 Real-Time PCR machine. qPCR primers used in this study are listed in Table 1.

### ELISAs

The following ELISA kits were used as directed by the manufacturer: Human IL-1β Set II (BD # 557953); Human Total IL-18 DuoSet ELISA (R&D Systems #DY318-05); ELISA MAX Standard Set Human TNFα (Biolegend #430201) or Human TNFα DuoSet ELISA (R&D Systems #DY210-05); ELISA MAX Standard Set Human IL-6 (Biolegend #430501); Human IL-8/CXCL8 DuoSet ELISA (R&D Systems #DY208-05).

### Luciferase Assay

The following plasmids were added to each well of a 96-well plate in 25ul of OptiMEM: 100ng of effector plasmid, 50ng of *Firefly* luciferase reporter plasmid (either NF-κB or ISRE Luc), and 2ng of *Renilla* luciferase plasmid. For assays using Pam3CSK4, 75ng of effector plasmid and 25ng of hTLR2 plasmid were used. For assays using increasing concentrations of effector-encoded vector, the indicated amounts were used. Lipofectamine 2000 Transfection Reagent was used as directed by the manufacturer. Briefly, 0.5ul of reagent was used per well in 25ul of OptiMEM and added to the plasmids in a 1:1 mix. This mix was incubated for 30mins at room temperature. HEK293T cells were resuspended in complete DMEM without phenol red and added to the plasmid mix at 5x10^4^ cells/well. Cells were incubated at 37°C and 5% CO2 overnight. The next day, cells were stimulated with rTNF or Pam3CSK4 for 4hrs, or rIL-1β or rIFNβ for 8hrs. The luciferase assay was conducted using the Dual Glo Luciferase Assay System (Promega #E2940) as directed by the manufacturer. Relative response ratios for the luciferase screen were calculated according to the manufacturer’s technical manual and using the following formula:

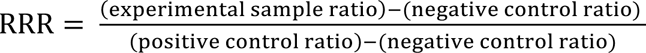

In this formula, the “ratio” refers to Firefly/Renilla ratio, the negative control refers to cells transfected with an mCherry-expressing vector, and the positive control refers to cells transfected with a YopJ-expressing vector, a *Yersinia* effector that potently inhibits NF-κB and MAPK signaling (Mukherjee et al., 2006; Orth et al., 2000; Zhou et al., 2005).

### Cell fractionation

For cell fractionation in HEK293Ts, cells were seeded at a concentration of 8x10^6^ cells/10cm dish and transfected overnight with 14ug of plasmid using Lipofectamine 2000 as directed by the manufacturer. For cell fractionation in THP-1s, cells were transduced as described above. Trypsin-EDTA 0.05% (Gibco, #25300054) was used to harvest HEK293T cells. Cells were counted and resuspended in NAR A Buffer (10mM HEPES pH 7.9, 10mM KCl, 0.1mM EDTA, 1mM DTT, supplemented with PMSF, NaF, and Protease and Phosphatase Inhibitor from Halt, #78440). Cells were allowed to swell on ice for 20 mins. NP-40 was added to a final concentration of 0.1% and samples were incubated for 1-1.5mins at room temperature.

Samples were vortexed gently for 15s and spun at 8500rpm at 4°C for 5 mins. The supernatant was transferred to a new tube and these cytoplasmic fractions were further spun at full speed at 4°C to remove any remaining cell debris. The nuclear fraction was washed 5x with NAR A buffer, with 1min spins at 8500rpm at 4°C in between. Chromatin fractions were obtained as previously described (Herrmann, Avgousti, & Weitzman, 2017). Briefly, the nuclei were resuspended in Buffer III.A and digested with MNase at 37°C for 30mins. The nuclei were further fragmented using buffers with increasing salt concentration (80-600mM NaCl) to extract the chromatin.

### Immunoprecipitation (IP) for mass spectrometry

HEK293T cells were seeded at 6x10^6^ cells/10cm dish, 2 plates per sample. The next day, cells were transfected with 14ug of either pgLAP1 empty vector (expressing GFP alone) or pgLAP1 encoding CBU1314 (expressing GFP-CBU1314). Each condition had 4 replicates. 24hrs later, cells were stimulated with 10ng/ml rTNF, harvested with trypsin, and crosslinked with paraformaldehyde in PBS at a final concentration (f.c.) of 1% for 20mins at room temperature. The PFA was quenched with glycine f.c. 125mM for 5mins at 37°C. Cells were washed once with ice-cold PBS and resuspended in 1ml of RIPA buffer (10 mM Tris/Cl pH 7.5, 150 mM NaCl, 0.5 mM EDTA, 0.1% SDS, 1% Triton X-100, 1% deoxycholate, filter sterilized) supplemented with protease inhibitors (Halt Protease and Phosphatase Inhibitor Cocktail, Roche Complete Mini EDTA-Free Protease Inhibitor Cocktail Tablets, PMSF f.c. 1mM). Each sample was sonicated on ice for 15 pulses, 30s on/30s off, at 25% amplitude using a Qsonica probe sonicator (Q55 model). MgCl2 was added to f.c. 2.5mM and samples were digested with rocking at room temperature for 30mins with DNase I (Roche #4716728001, 150U/ml). NaCl was added to f.c. 600mM and samples were incubated at 4°C with rotation for 20mins to extract chromatin proteins. Samples were centrifuged at 20,000xg at 4°C for 10mins, and the supernatants were diluted by adding 2ml of Wash Buffer II (10mM Tris-HCl pH 7.5, 150mM NaCl) supplemented with protease inhibitors. GFP-Trap magnetic beads (Chromotek #gtd-100) were washed 3X with Wash Buffer I (10mM Tris-HCl pH 7.5, 150mM NaCl, 0.25% NP-40), 200ul beads/sample, and added to each sample. Samples were rotated end over end for 30mins at 4°C. Beads were separated with a magnet and washed 1X with Wash Buffer I, then 3X with Wash Buffer II. On the last wash, samples were transferred to new tubes. Beads were submitted for mass spectrometry analysis to The Children’s Hospital of Philadelphia (CHOP) Proteomics Core.

### Mass spectrometry (MS) analysis

#### In solution digestion

Proteins were solubilized and digested with the iST kit (PreOmics GmbH, Martinsried, Germany) per manufacturer’s protocol. Briefly, the proteins were solubilized, reduced and alkylated by addition of SDC buffer containing TCEP and 2-chloroacetamide to the washed beads then heated to 95°C for 10mins. Proteins were enzymatically hydrolyzed for 1.5 hours at 37°C by addition of LysC and trypsin. The resulting peptides were de-salted, dried by vacuum centrifugation, and reconstituted in 0.1% TFA containing iRT peptides (Biognosys, Schlieren, Switzerland).

#### MS data acquisition

Samples were analyzed on an Exploris 480 mass spectrometer (Thermo Fisher Scientific San Jose, CA) coupled with an Ultimate 3000 nano UPLC system and an EasySpray source. Peptides were loaded onto an Acclaim PepMap 100 75um x 2cm trap column (Thermo) at 5uL/min, and separated by reverse phase (RP)-HPLC on a nanocapillary column, 75 μm id × 50cm 2um PepMap RSLC C18 column (Thermo). Mobile phase A consisted of 0.1% formic acid and mobile phase B of 0.1% formic acid/acetonitrile. Peptides were eluted into the mass spectrometer at 300 nL/min with each RP-LC run comprising a 90 minute gradient from 3% B to 38% B. The mass spectrometer was set with a master scan at R=120000, a scan range of 300-1400, AGC target set to standard, maximum injection time set to auto, and dynamic exclusion set to 30 seconds repeat 1. Charge state 2-5 were included. Top 15 data dependent MSMS scans were collected at R=45000, first mass set to 120, normalized AGC target at 300%, maximum injection time=auto, HCD NCE set to 30.

#### MS raw data processing

Peptide and protein identification/quantification was performed with MaxQuant (2.0.3.0) (Tyanova, Temu, & Cox, 2016) using a human reference database from Uniprot (reviewed canonical and isoforms) appended with the sequence of iRT peptides.

Carbamidomethyl of Cys was defined as a fixed modification. Oxidation of Met and acetylation of protein N-terminal were set as variable modifications. Trypsin/P was selected as the digestion enzyme, and a maximum of 3 labeled amino acids and 2 missed cleavages per peptide were allowed. The False Discovery Rate for peptides and proteins was set at 1%. Fragment ion tolerance was set to 0.5 Da. The MS/MS tolerance was set at 20 ppm. The minimum peptide length was set at 7 amino acids. The rest of the parameters were kept as default. The quality of the generated results by MaxQuant was further visualized and verified by PTXQC (Bielow, Mastrobuoni, & Kempa, 2016).

#### Data processing and statistical analysis

Perseus (1.6.14.0) (Tyanova, Temu, Sinitcyn, et al., 2016) was used for proteomics data processing and statistical analysis. The MaxLFQ intensity values were used to analyze the whole cell proteome data. Protein groups containing matches to decoy database or contaminants were discarded. The data were log2 transformed and normalized by subtracting the median for each sample. Proteins with less than 4 valid values in at least one group were filtered out. Student’s t-test was employed to identify differentially expressed proteins and volcano plots were generated to visualize the affected proteins while comparing different groups of samples. We used Benjamini-Hochberg FDR of 0.05 as false discovery control (i.e. significant hits should have a FDR of less than 0.05). The proteins that were exclusively detected in one experimental group were also reported for further bioinformatics analysis.

### Co-IP experiments

Co-IPs were conducted as described above for IP, with the following changes: one 10-cm dish/sample, sonicated for 10 pulses; 50ul of GFP-Trap beads/sample, rotated at 4°C for 45min; beads were washed 3X with Wash Buffer II only; protein was eluted from beads by resuspending in 4X Laemmli Buffer (Bio-Rad #1610747) with 100mM DTT, and boiled for 20mins to reverse crosslinks.

### Western blots

Protein samples were quantified using the Bio-Rad Protein Assay Kit II as indicated by the manufacturer (Bio-Rad, #5000002), and normalized before running the immunoblots. For co-immunoprecipitation experiments, the samples were quantified and normalized prior to the IP. Samples were run on 12% SDS-PAGE gels and transferred to PVDF membranes using the Mini-PROTEAN Tetra Cell System (Bio-Rad). For large targets, samples were run on a 6% gel. Membranes were blocked overnight at 4°C with 5% milk. The next day, membranes were incubated with the indicated primary antibodies for 2hrs at room temperature, washed 3x for 10 mins with TBS-T Buffer (0.2M Tris base, 1.37M NaCl, 0.1% Tween-20), and incubated with HRP-conjugated secondary antibody for 1hr at room temperature. Membranes were washed 3x for 10 mins with TBS-T Buffer, and developed with ECL Western Blotting Substrate (Pierce, #32106) or SuperSignal West Femto Maximum Sensitivity Substrate (Pierce, #34095). Where indicated, Western blot bands were quantified using Image J.

The following antibodies were used in this study: p-IKKα/β (CST 2697S, 1:1000), IKKα (Novus Biologicals NB 100-56704SS, 1:1000), p-p65 RelA (CST 3033S, 1:1000), p65 RelA (CST 8242S, 1:1000), IκBα (CST 4814T, 1:1000), p38 (CST 9212, 1:1000), p-SAPK/JNK (CST 9251S, 1:1000), SAPK/JNK (CST 9258P, 1:1000), p-ERK1/2 (CST 4370P, 1:1000), ERK1/2 (CST 4695T, 1:1000), β-actin (CST 4967L, 1:1000), Streptag-HRP (EMD Millipore 71591-75, 1:4000), GFP (CST 2956, 1:1000), α-Tubulin (Invitrogen A11126, 1:5000), H3 (CST 4620S, 1:1000), IRG1 (CST 77510S, 1:1000), IL-1β (R&D Systems MAB201-100, 1:500), CDC73 (CST 8126S, 1:1000), PAF1 (CST 12883S, 1:1000), CTR9 (CST 12619S, 1:1000), RPB1-CTD (2629S, 1:1000), Anti-Rabbit IgG HRP-linked Antibody (CST 7074S, 1:2000), and Anti-Mouse IgG HRP-linked Antibody (CST 7076S, 1:2000).

### siRNA knockdown with luciferase assay in HEK293T cells

Lipofectamine RNAiMAX Transfection Reagent (Invitrogen) was used to transfect 2pmol of control siRNA or siRNA specific for genes of interest per well of a 96-well plate, following the manufacturer’s instructions. siRNA was added to cells at the time of seeding, as mentioned above for the luciferase assay. The next day, cells were transfected as described above with the luciferase reporter plasmids. 24hrs later, cells were stimulated with the indicated agonists and the luciferase assay was conducted as previously mentioned. The following Silencer Select siRNAs (Life Technologies) were used: Control (non-targeting) siRNAs: Negative Control No. 1 (#4390843) and No. 2 (#4390846) Human *PAF1* siRNA IDs: s29267 and s29268 Human *CDC73* siRNA ID: s35849 Human *AKAP8* siRNA ID: s20070 Human *HSPA6/7* siRNA ID: s6983 Human *MYO5C* siRNA ID: s31794

### siRNA knockdown in HEK293T cells for growth curves

HEK293T cells were seeded at 1x10^5^ cells/well in a 24-well plate, and transfected with 12pmol of siRNA using Lipofectamine RNAiMAX, following the manufacturer’s instructions. The next day, the media was replaced to remove the Lipofectamine. 48hrs post-siRNA transfection, cells were infected with mCherry-expressing *C. burnetii* at an MOI of 100. 4hrs post-infection, cells were washed with PBS and day 0 samples were harvested with water. The media for day 7 samples was replaced after washing. On day 3, cells were fed with 500ul of media. On day 4, cells were harvested for GE analysis. The siRNAs used are listed in the above section.

### siRNA knockdown in murine BMDMs for mRNA expression and growth curves

BMDMs were replated into 24-well plates at 1.5x10^5^ cells/well and transfected with 12pmol of siRNA using Lipofectamine RNAiMAX, following the manufacturer’s instructions. The next day, the media was replaced to remove the Lipofectamine. <u>For mRNA expression</u>: 48-hrs post-siRNA transfection, cells were infected with WT or *icmL*::Tn *C. burnetii* at MOI 50 and RNA was harvested for analysis at 24hrs post-infection. The murine primers in Table 1 were used for gene expression analysis. <u>For growth curves</u>: 48hrs post-siRNA transfection, cells were infected with mCherry-expressing *C. burnetii* at an MOI of 100. 4hrs post-infection, cells were washed with PBS and day 0 samples were harvested with water. The media for day 7 samples was replaced after washing. On day 3, cells were transfected with another round of siRNA, and harvested on day 7 for GE analysis. The control siRNAs were the same as for the human cells experiments. The following Silencer Select siRNAs were used for murine *Paf1*: Murine *Paf1* siRNA IDs: s79773 and s79774.

### siRNA knockdown in THP-1 monocytes for mRNA expression

Lipofectamine RNAiMAX transfection reagent (Invitrogen) was used to transfect 12pmol of control or *PAF1* siRNA per well of a 24-well plate, following the manufacturer’s instructions. siRNA was added to cells at the time of seeding, with 2x10^5^ cells/well in replating media. The next day, 0.4ml of replating media was added to each well to dilute the transfection reagent. 48hrs post-siRNA transfection, cells were stimulated for 6hrs with TNF (1ng/ml) or IFNβ (1U/ml) and harvested for mRNA expression analysis. Alternatively, cells were infected with WT or *icmL*::Tn *C. burnetii* prior to RNA harvest.

### Statistical Analysis

All graphs and statistical analyses were generated using Prism (GraphPad Software). Statistical analyses were carried out as indicated in each figure legend. For all analyses: *p≤0.05; **p≤0.01; ***p≤0.001; ****p≤0.0001.

## Acknowledgements

We thank members of the laboratories of Sunny Shin and Igor Brodsky at the University of Pennsylvania, Michael May at the University of Pennsylvania, Matthew Weitzman and members of his laboratory at the Children’s Hospital of Philadelphia, and James Samuel, Erin van Schaik, and their laboratory members at Texas A&M College of Medicine for scientific discussions and advice. We thank Igor Brodsky for critical reading of the manuscript. We thank Paul Bates and Kellie Jurado at the University of Pennsylvania for reagents and protocols. We thank the Human Immunology Core of the Penn Center for AIDS Research and the Abramson Cancer Center, which is supported by NIH/NCI grant P30-CA016520, for providing purified primary human monocytes. The Shin laboratory is supported by NIH/NIAID grants AI118861, AI123243, and AI161476 and the Linda Pechenik Montague Investigator Award from the University of Pennsylvania Perelman School of Medicine. S.S. is a recipient of the Burroughs-Wellcome Fund Investigators in the Pathogenesis of Infectious Disease Award. N.L.F. is a recipient of the NIH Cell and Molecular Biology T32GM07229 and the HHMI James H. Gilliam, Jr. Fellowship for Advanced Study.

## References

Abdel-Mohsen, M., Wang, C., Strain, M. C., Lada, S. M., Deng, X., Cockerham, L. R., . . . Pillai, S. K. (2015). Select host restriction factors are associated with HIV persistence during antiretroviral therapy. AIDS, 29(4), 411–420. doi:10.1097/QAD.0000000000000572

Ammerdorffer, A., Schoffelen, T., Gresnigt, M. S., Oosting, M., den Brok, M. H., Abdollahi-Roodsaz, S., . . . Sprong, T. (2015). Recognition of Coxiella burnetii by toll-like receptors and nucleotide-binding oligomerization domain-like receptors. J Infect Dis, 211(6), 978–987. doi:10.1093/infdis/jiu526

Beare, P. A. (2012). Genetic manipulation of Coxiella burnetii. Adv Exp Med Biol, 984, 249–271. doi:10.1007/978-94-007-4315-1_13

Beare, P. A., Howe, D., Cockrell, D. C., Omsland, A., Hansen, B., & Heinzen, R. A. (2009). Characterization of a Coxiella burnetii ftsZ mutant generated by Himar1 transposon mutagenesis. J Bacteriol, 191(5), 1369–1381. doi:10.1128/JB.01580-08

Bielow, C., Mastrobuoni, G., & Kempa, S. (2016). Proteomics Quality Control: Quality Control Software for MaxQuant Results. J Proteome Res, 15(3), 777–787. doi:10.1021/acs.jproteome.5b00780

Bisle, S., Klingenbeck, L., Borges, V., Sobotta, K., Schulze-Luehrmann, J., Menge, C., . . . Lührmann, A. (2016). The inhibition of the apoptosis pathway by the Coxiella burnetii effector protein CaeA requires the EK repetition motif, but is independent of survivin. Virulence, 7(4), 400–412. doi:10.1080/21505594.2016.1139280

Boucherit, N., Barry, A. O., Mottola, G., Trouplin, V., Capo, C., Mege, J. L., & Ghigo, E. (2012). Effects of Coxiella burnetii on MAPKinases phosphorylation. FEMS Immunol Med Microbiol, 64(1), 101–103. doi:10.1111/j.1574-695X.2011.00852.x

Bradley, W. P., Boyer, M. A., Nguyen, H. T., Birdwell, L. D., Yu, J., Ribeiro, J. M., . . . Shin, S. (2016). Primary Role for Toll-Like Receptor-Driven Tumor Necrosis Factor Rather than Cytosolic Immune Detection in Restricting Coxiella burnetii Phase II Replication within Mouse Macrophages. Infect Immun, 84(4), 998–1015. doi:10.1128/IAI.01536-15

Burette, M., Allombert, J., Lambou, K., Maarifi, G., Nisole, S., Di Russo Case, E., . . . Bonazzi, M. (2020). Modulation of innate immune signaling by a *Coxiella burnetii* eukaryotic-like effector protein. Proc Natl Acad Sci U S A, 117(24), 13708–13718. doi:10.1073/pnas.1914892117

Carey, K. L., Newton, H. J., Lührmann, A., & Roy, C. R. (2011). The Coxiella burnetii Dot/Icm system delivers a unique repertoire of type IV effectors into host cells and is required for intracellular replication. PLoS Pathog, 7(5), e1002056. doi:10.1371/journal.ppat.1002056

Chen, C., Banga, S., Mertens, K., Weber, M. M., Gorbaslieva, I., Tan, Y., . . . Samuel, J. E. (2010). Large-scale identification and translocation of type IV secretion substrates by Coxiella burnetii. Proc Natl Acad Sci U S A, 107(50), 21755–21760. doi:10.1073/pnas.1010485107

Cordsmeier, A., Rinkel, S., Jeninga, M., Schulze-Luehrmann, J., Ölke, M., Schmid, B., . . . Lührmann, A. (2022). The Coxiella burnetii T4SS effector protein AnkG hijacks the 7SK small nuclear ribonucleoprotein complex for reprogramming host cell transcription. PLoS Pathog, 18(2), e1010266. doi:10.1371/journal.ppat.1010266

Cunha, L. D., Ribeiro, J. M., Fernandes, T. D., Massis, L. M., Khoo, C. A., Moffatt, J. H., . . . Zamboni, D. S. (2015). Inhibition of inflammasome activation by Coxiella burnetii type IV secretion system effector IcaA. Nat Commun, 6, 10205. doi:10.1038/ncomms10205

DeLaney, A. A., Berry, C. T., Christian, D. A., Hart, A., Bjanes, E., Wynosky-Dolfi, M. A., . . . Brodsky, I. E. (2019). Caspase-8 promotes c-Rel-dependent inflammatory cytokine expression and resistance against. Proc Natl Acad Sci U S A, 116(24), 11926–11935. doi:10.1073/pnas.1820529116

Dorrington, M. G., & Fraser, I. D. C. (2019). NF-κB Signaling in Macrophages: Dynamics, Crosstalk, and Signal Integration. Front Immunol, 10, 705. doi:10.3389/fimmu.2019.00705

Dragan, A. L., Kurten, R. C., & Voth, D. E. (2019). Characterization of Early Stages of Human Alveolar Infection by the Q Fever Agent. Infect Immun, 87(5). doi:10.1128/IAI.00028-19

Finlay, B. B., & McFadden, G. (2006). Anti-immunology: evasion of the host immune system by bacterial and viral pathogens. Cell, 124(4), 767–782. doi:10.1016/j.cell.2006.01.034

Francette, A. M., Tripplehorn, S. A., & Arndt, K. M. (2021). The Paf1 Complex: A Keystone of Nuclear Regulation Operating at the Interface of Transcription and Chromatin. J Mol Biol, 433(14), 166979. doi:10.1016/j.jmb.2021.166979

Graham, J. G., MacDonald, L. J., Hussain, S. K., Sharma, U. M., Kurten, R. C., & Voth, D. E. (2013). Virulent Coxiella burnetii pathotypes productively infect primary human alveolar macrophages. Cell Microbiol, 15(6), 1012–1025. doi:10.1111/cmi.12096

Graham, J. G., Winchell, C. G., Kurten, R. C., & Voth, D. E. (2016). Development of an Ex Vivo Tissue Platform To Study the Human Lung Response to Coxiella burnetii. Infect Immun, 84(5), 1438–1445. doi:10.1128/IAI.00012-16

Herrmann, C., Avgousti, D. C., & Weitzman, M. D. (2017). Differential Salt Fractionation of Nuclei to Analyze Chromatin-associated Proteins from Cultured Mammalian Cells. Bio Protoc, 7(6). doi:10.21769/BioProtoc.2175

Hou, L., Wang, Y., Liu, Y., Zhang, N., Shamovsky, I., Nudler, E., . . . Dynlacht, B. D. (2019). Paf1C regulates RNA polymerase II progression by modulating elongation rate. Proc Natl Acad Sci U S A, 116(29), 14583–14592. doi:10.1073/pnas.1904324116

Janeway, C. A., & Medzhitov, R. (2002). Innate immune recognition. Annu Rev Immunol, 20, 197–216. doi:10.1146/annurev.immunol.20.083001.084359

Kak, G., Raza, M., & Tiwari, B. K. (2018). Interferon-gamma (IFN-γ): Exploring its implications in infectious diseases. Biomol Concepts, 9(1), 64–79. doi:10.1515/bmc-2018-0007

Kawai, T., & Akira, S. (2011). Toll-like receptors and their crosstalk with other innate receptors in infection and immunity. Immunity, 34(5), 637–650. doi:10.1016/j.immuni.2011.05.006

Kenaston, M. W., Pham, O. H., Petit, M. J., & Shah, P. S. (2022). PAF1 modulates innate immunity by both activation and repression of gene expression. In: BioRXIV.

Kim, J., Guermah, M., & Roeder, R. G. (2010). The human PAF1 complex acts in chromatin transcription elongation both independently and cooperatively with SII/TFIIS. Cell, 140(4), 491–503. doi:10.1016/j.cell.2009.12.050

Klingenbeck, L., Eckart, R. A., Berens, C., & Lührmann, A. (2013). The Coxiella burnetii type IV secretion system substrate CaeB inhibits intrinsic apoptosis at the mitochondrial level. Cell Microbiol, 15(4), 675–687. doi:10.1111/cmi.12066

Kopp, E. B., & Ghosh, S. (1995). NF-kappa B and rel proteins in innate immunity. Adv Immunol, 58, 1–27. doi:10.1016/s0065-2776(08)60618-5

Larson, C. L., Martinez, E., Beare, P. A., Jeffrey, B., Heinzen, R. A., & Bonazzi, M. (2016). Right on Q: genetics begin to unravel Coxiella burnetii host cell interactions. Future Microbiol, 11, 919–939. doi:10.2217/fmb-2016-0044

Lifshitz, Z., Burstein, D., Schwartz, K., Shuman, H. A., Pupko, T., & Segal, G. (2014). Identification of novel Coxiella burnetii Icm/Dot effectors and genetic analysis of their involvement in modulating a mitogen-activated protein kinase pathway. Infect Immun, 82(9), 3740–3752. doi:10.1128/IAI.01729-14

Liu, L., Oliveira, N. M., Cheney, K. M., Pade, C., Dreja, H., Bergin, A. M., . . . McKnight, A. (2011). A whole genome screen for HIV restriction factors. Retrovirology, 8, 94. doi:10.1186/1742-4690-8-94

Lopes Fischer, N., Naseer, N., Shin, S., & Brodsky, I. E. (2020). Effector-triggered immunity and pathogen sensing in metazoans. Nat Microbiol, 5(1), 14–26. doi:10.1038/s41564-019-0623-2

Luan, H. H., & Medzhitov, R. (2016). Food Fight: Role of Itaconate and Other Metabolites in Antimicrobial Defense. Cell Metab, 24(3), 379–387. doi:10.1016/j.cmet.2016.08.013

Lührmann, A., Nogueira, C. V., Carey, K. L., & Roy, C. R. (2010). Inhibition of pathogen-induced apoptosis by a Coxiella burnetii type IV effector protein. Proc Natl Acad Sci U S A, 107(44), 18997–19001. doi:10.1073/pnas.1004380107

Madariaga, M. G., Rezai, K., Trenholme, G. M., & Weinstein, R. A. (2003). Q fever: a biological weapon in your backyard. Lancet Infect Dis, 3(11), 709–721.

Mahapatra, S., Gallaher, B., Smith, S. C., Graham, J. G., Voth, D. E., & Shaw, E. I. (2016). *Coxiella burnetii* Employs the Dot/Icm Type IV Secretion System to Modulate Host NF- κB/RelA Activation. Front Cell Infect Microbiol, 6, 188. doi:10.3389/fcimb.2016.00188

Marazzi, I., Ho, J. S., Kim, J., Manicassamy, B., Dewell, S., Albrecht, R. A., . . . Tarakhovsky, A. (2012). Suppression of the antiviral response by an influenza histone mimic. Nature, 483(7390), 428–433. doi:10.1038/nature10892

Maurin, M., & Raoult, D. (1999). Q fever. Clin Microbiol Rev, 12(4), 518–553.

Medzhitov, R. (2007). Recognition of microorganisms and activation of the immune response. Nature, 449(7164), 819–826. doi:10.1038/nature06246

Meghari, S., Honstettre, A., Lepidi, H., Ryffel, B., Raoult, D., & Mege, J. L. (2005). TLR2 is necessary to inflammatory response in Coxiella burnetii infection. Ann N Y Acad Sci, 1063, 161–166. doi:10.1196/annals.1355.025

Michelucci, A., Cordes, T., Ghelfi, J., Pailot, A., Reiling, N., Goldmann, O., . . . Hiller, K. (2013). Immune-responsive gene 1 protein links metabolism to immunity by catalyzing itaconic acid production. Proc Natl Acad Sci U S A, 110(19), 7820–7825. doi:10.1073/pnas.1218599110

Mitchell, S., Vargas, J., & Hoffmann, A. (2016). Signaling via the NFκB system. Wiley Interdiscip Rev Syst Biol Med, 8(3), 227–241. doi:10.1002/wsbm.1331

Moffatt, J. H., Newton, P., & Newton, H. J. (2015). Coxiella burnetii: turning hostility into a home. Cell Microbiol, 17(5), 621–631. doi:10.1111/cmi.12432

Mukherjee, S., Keitany, G., Li, Y., Wang, Y., Ball, H. L., Goldsmith, E. J., & Orth, K. (2006). Yersinia YopJ acetylates and inhibits kinase activation by blocking phosphorylation. Science, 312(5777), 1211–1214. doi:10.1126/science.1126867

Naujoks, J., Tabeling, C., Dill, B. D., Hoffmann, C., Brown, A. S., Kunze, M., . . . Opitz, B. (2016). IFNs Modify the Proteome of Legionella-Containing Vacuoles and Restrict Infection Via IRG1-Derived Itaconic Acid. PLoS Pathog, 12(2), e1005408. doi:10.1371/journal.ppat.1005408

Omsland, A., Cockrell, D. C., Howe, D., Fischer, E. R., Virtaneva, K., Sturdevant, D. E., . . . Heinzen, R. A. (2009). Host cell-free growth of the Q fever bacterium Coxiella burnetii. Proc Natl Acad Sci U S A, 106(11), 4430–4434. doi:10.1073/pnas.0812074106

Omsland, A., & Heinzen, R. A. (2011). Life on the outside: the rescue of Coxiella burnetii from its host cell. Annu Rev Microbiol, 65, 111–128. doi:10.1146/annurev-micro-090110-102927

Orth, K., Xu, Z., Mudgett, M. B., Bao, Z. Q., Palmer, L. E., Bliska, J. B., . . . Dixon, J. E. (2000). Disruption of signaling by Yersinia effector YopJ, a ubiquitin-like protein protease. Science, 290(5496), 1594–1597.

Palazon, A., Goldrath, A. W., Nizet, V., & Johnson, R. S. (2014). HIF transcription factors, inflammation, and immunity. Immunity, 41(4), 518–528. doi:10.1016/j.immuni.2014.09.008

Parnas, O., Jovanovic, M., Eisenhaure, T. M., Herbst, R. H., Dixit, A., Ye, C. J., . . . Regev, A. (2015). A Genome-wide CRISPR Screen in Primary Immune Cells to Dissect Regulatory Networks. Cell, 162(3), 675–686. doi:10.1016/j.cell.2015.06.059

Patrick, K. L., Wojcechowskyj, J. A., Bell, S. L., Riba, M. N., Jing, T., Talmage, S., . . . Watson, R. O. (2018). Quantitative Yeast Genetic Interaction Profiling of Bacterial Effector Proteins Uncovers a Role for the Human Retromer in Salmonella Infection. Cell Syst, 7(3), 323–338.e326. doi:10.1016/j.cels.2018.06.010

Petit, M. J., Kenaston, M. W., Pham, O. H., Nagainis, A. A., Fishburn, A. T., & Shah, P. S. (2021). Nuclear dengue virus NS5 antagonizes expression of PAF1-dependent immune response genes. PLoS Pathog, 17(11), e1010100. doi:10.1371/journal.ppat.1010100

Platanias, L. C. (2005). Mechanisms of type-I- and type-II-interferon-mediated signalling. Nat Rev Immunol, 5(5), 375–386. doi:10.1038/nri1604

Rahman, M. M., & McFadden, G. (2011). Modulation of NF-κB signalling by microbial pathogens. Nat Rev Microbiol, 9(4), 291–306. doi:10.1038/nrmicro2539

Ramstead, A. G., Robison, A., Blackwell, A., Jerome, M., Freedman, B., Lubick, K. J., . . . Jutila, M. A. (2016). Roles of Toll-Like Receptor 2 (TLR2), TLR4, and MyD88 during Pulmonary Coxiella burnetii Infection. Infect Immun, 84(4), 940–949. doi:10.1128/IAI.00898-15

Raposo, R. A., Abdel-Mohsen, M., Bilska, M., Montefiori, D. C., Nixon, D. F., & Pillai, S. K. (2013). Effects of cellular activation on anti-HIV-1 restriction factor expression profile in primary cells. J Virol, 87(21), 11924–11929. doi:10.1128/JVI.02128-13

Rotz, L. D., Khan, A. S., Lillibridge, S. R., Ostroff, S. M., & Hughes, J. M. (2002). Public health assessment of potential biological terrorism agents. Emerg Infect Dis, 8(2), 225–230. doi:10.3201/eid0802.010164

Roy, C. R., & Mocarski, E. S. (2007). Pathogen subversion of cell-intrinsic innate immunity. Nat Immunol, 8(11), 1179–1187. doi:10.1038/ni1528

Schneeberger, P. M., Wintenberger, C., van der Hoek, W., & Stahl, J. P. (2014). Q fever in the Netherlands - 2007-2010: what we learned from the largest outbreak ever. Med Mal Infect, 44(8), 339–353. doi:10.1016/j.medmal.2014.02.006

Shah, P. S., Link, N., Jang, G. M., Sharp, P. P., Zhu, T., Swaney, D. L., . . . Krogan, N. J. (2018). Comparative Flavivirus-Host Protein Interaction Mapping Reveals Mechanisms of Dengue and Zika Virus Pathogenesis. Cell, 175(7), 1931–1945.e1918. doi:10.1016/j.cell.2018.11.028

Smale, S. T. (2012). Transcriptional regulation in the innate immune system. Curr Opin Immunol, 24(1), 51–57. doi:10.1016/j.coi.2011.12.008

Tyanova, S., Temu, T., & Cox, J. (2016). The MaxQuant computational platform for mass spectrometry-based shotgun proteomics. Nat Protoc, 11(12), 2301–2319. doi:10.1038/nprot.2016.136

Tyanova, S., Temu, T., Sinitcyn, P., Carlson, A., Hein, M. Y., Geiger, T., . . . Cox, J. (2016). The Perseus computational platform for comprehensive analysis of (prote)omics data. Nat Methods, 13(9), 731–740. doi:10.1038/nmeth.3901

Viatour, P., Merville, M. P., Bours, V., & Chariot, A. (2005). Phosphorylation of NF-kappaB and IkappaB proteins: implications in cancer and inflammation. Trends Biochem Sci, 30(1), 43–52. doi:10.1016/j.tibs.2004.11.009

Voth, D. E., & Heinzen, R. A. (2009). Sustained activation of Akt and Erk1/2 is required for Coxiella burnetii antiapoptotic activity. Infect Immun, 77(1), 205–213. doi:10.1128/IAI.01124-08

Weber, M. M., Faris, R., McLachlan, J., Tellez, A., Wright, W. U., Galvan, G., . . . Samuel, J. E. (2016). Modulation of the host transcriptome by Coxiella burnetii nuclear effector Cbu1314. Microbes Infect, 18(5), 336–345. doi:10.1016/j.micinf.2016.01.003

Wuerth, J. D., & Weber, F. (2016). Phleboviruses and the Type I Interferon Response. Viruses, 8(6). doi:10.3390/v8060174

Yang, Y., Li, W., Hoque, M., Hou, L., Shen, S., Tian, B., & Dynlacht, B. D. (2016). PAF Complex Plays Novel Subunit-Specific Roles in Alternative Cleavage and Polyadenylation. PLoS Genet, 12(1), e1005794. doi:10.1371/journal.pgen.1005794

Youn, M. Y., Yoo, H. S., Kim, M. J., Hwang, S. Y., Choi, Y., Desiderio, S. V., & Yoo, J. Y. (2007). hCTR9, a component of Paf1 complex, participates in the transcription of interleukin 6-responsive genes through regulation of STAT3-DNA interactions. J Biol Chem, 282(48), 34727–34734. doi:10.1074/jbc.M705411200

Yu, M., Yang, W., Ni, T., Tang, Z., Nakadai, T., Zhu, J., & Roeder, R. G. (2015). RNA polymerase II-associated factor 1 regulates the release and phosphorylation of paused RNA polymerase II. Science, 350(6266), 1383–1386. doi:10.1126/science.aad2338

Zamboni, D. S., Campos, M. A., Torrecilhas, A. C., Kiss, K., Samuel, J. E., Golenbock, D. T., . . . Gazzinelli, R. T. (2004). Stimulation of toll-like receptor 2 by Coxiella burnetii is required for macrophage production of pro-inflammatory cytokines and resistance to infection. J Biol Chem, 279(52), 54405–54415. doi:10.1074/jbc.M410340200

Zhang, W., & Liu, H. T. (2002). MAPK signal pathways in the regulation of cell proliferation in mammalian cells. Cell Res, 12(1), 9–18. doi:10.1038/sj.cr.7290105

Zhang, Y., Fu, J., Liu, S., Wang, L., Qiu, J., van Schaik, E. J., . . . Luo, Z. Q. (2022). *Coxiella burnetii* inhibits host immunity by a protein phosphatase adapted from glycolysis. Proc Natl Acad Sci U S A, 119(1). doi:10.1073/pnas.2110877119

Zhou, H., Monack, D. M., Kayagaki, N., Wertz, I., Yin, J., Wolf, B., & Dixit, V. M. (2005). Yersinia virulence factor YopJ acts as a deubiquitinase to inhibit NF-kappa B activation. J Exp Med, 202(10), 1327–1332. doi:10.1084/jem.20051194

