## Supplemental Figures for "A *Coxiella burnetii* effector interacts with the host PAF1 complex and suppresses the innate immune response"

### Supplemental Figure 1

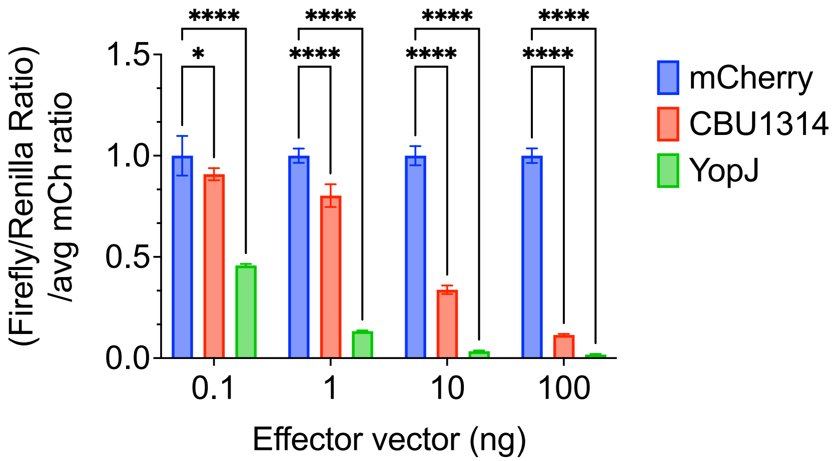

**Supplemental Figure 1. *C. burnetii* effector CBU1314 suppresses TNF-induced NF-κB signaling in a dose-dependent manner.** HEK293T cells were transfected with each effector-containing vector (pgLAP1) and the luciferase reporter system. Increasing amounts of effector vector were used, as indicated. 24hrs later, cells were stimulated with TNF at 1ng/ml for 4hrs prior to the luciferase assay. Firefly/Renilla ratios were further divided by the mCherry control ratio (mCh) for each concentration to account for differences in vector uptake due to increasing concentrations of DNA. Statistical analysis: Two-way ANOVA with Dunnett's multiple comparisons test within each concentration was conducted.

### Supplemental Figure 2

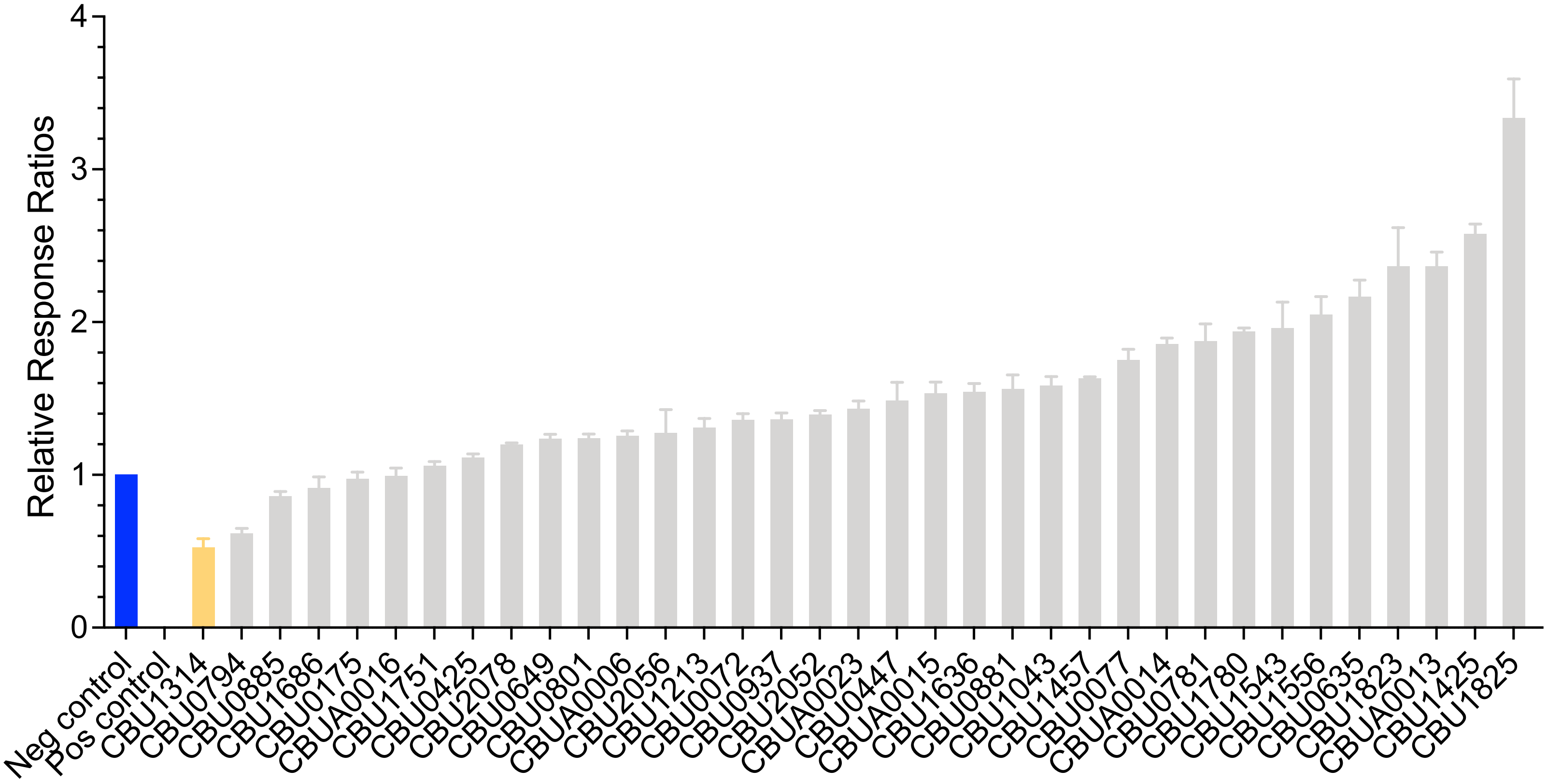

**Supplemental Figure 2. NF-κB luciferase screen using IL-1β stimulation.** HEK293T cells were transfected with each effector-containing vector (pgLAP1) and the luciferase reporter system. 24hrs post transfection, cells were stimulated using 0.1ng/ml of IL-1β. Luciferase signals were measured 8hrs post stimulation.

### Supplemental Figure 3

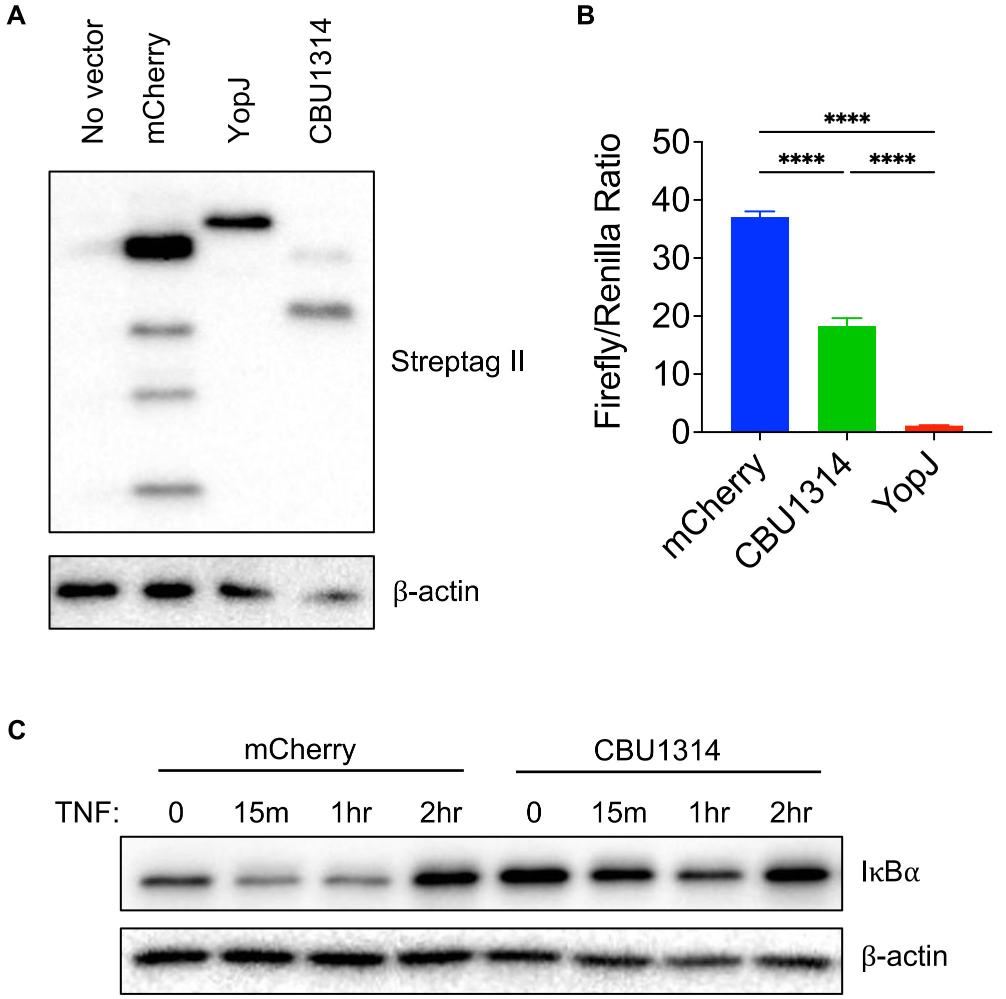

**Supplemental Figure 3. A C-terminally tagged CBU1314 in lentivector background also suppresses NF-κB signaling in HEK293T cells.** HEK293T cells were transfected with lentivectors encoding mCherry, CBU1314, or YopJ that were C-terminally tagged with a strep tag. A) Immunoblot analysis with antibodies against the strep tag showing expression of each vector as indicated. B) Cells were additionally transfected with the NF-κB luciferase reporter system. 24hrs post transfection, cells were stimulated with TNF at 1ng/ml for 4 hrs prior to conducting the luciferase assay. Statistical analysis: One-way ANOVA with multiple comparisons. C) Immunoblot for IκBα degradation during expression of mCherry or CBU1314 with a time course of TNF stimulation at 10ng/mL.

### Supplemental Figure 4

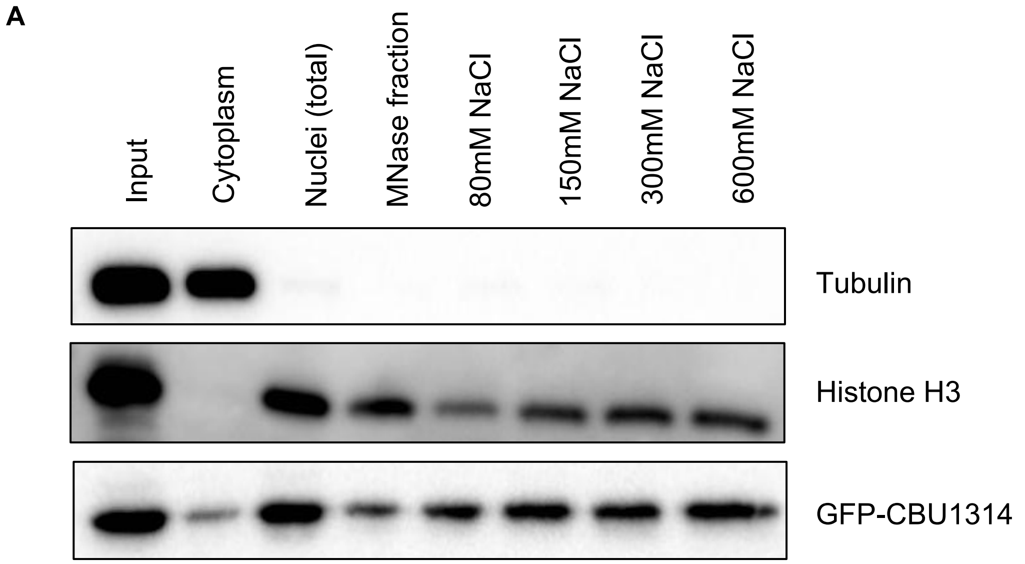


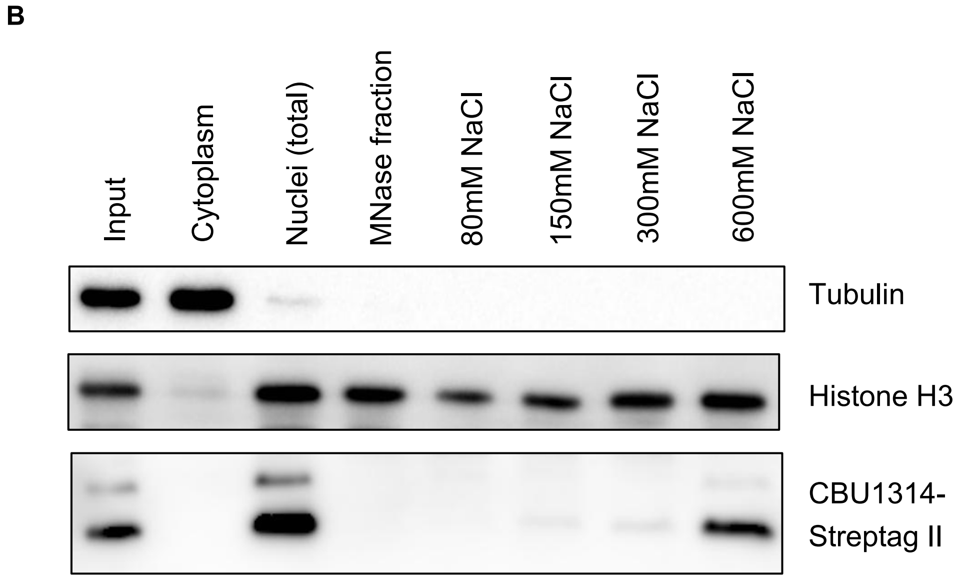

**Supplemental Figure 4. CBU1314 predominantly localizes to the nucleus and associates with chromatin.** A) HEK293T cells were transfected with a GFP-tagged CBU1314, and B) THP-1 cells were transduced with a strep-tagged CBU1314. 24hrs post transfection, cells were harvested for cell fractionation experiments using increasing salt concentrations for chromatin extraction.

### Supplemental Figure 5

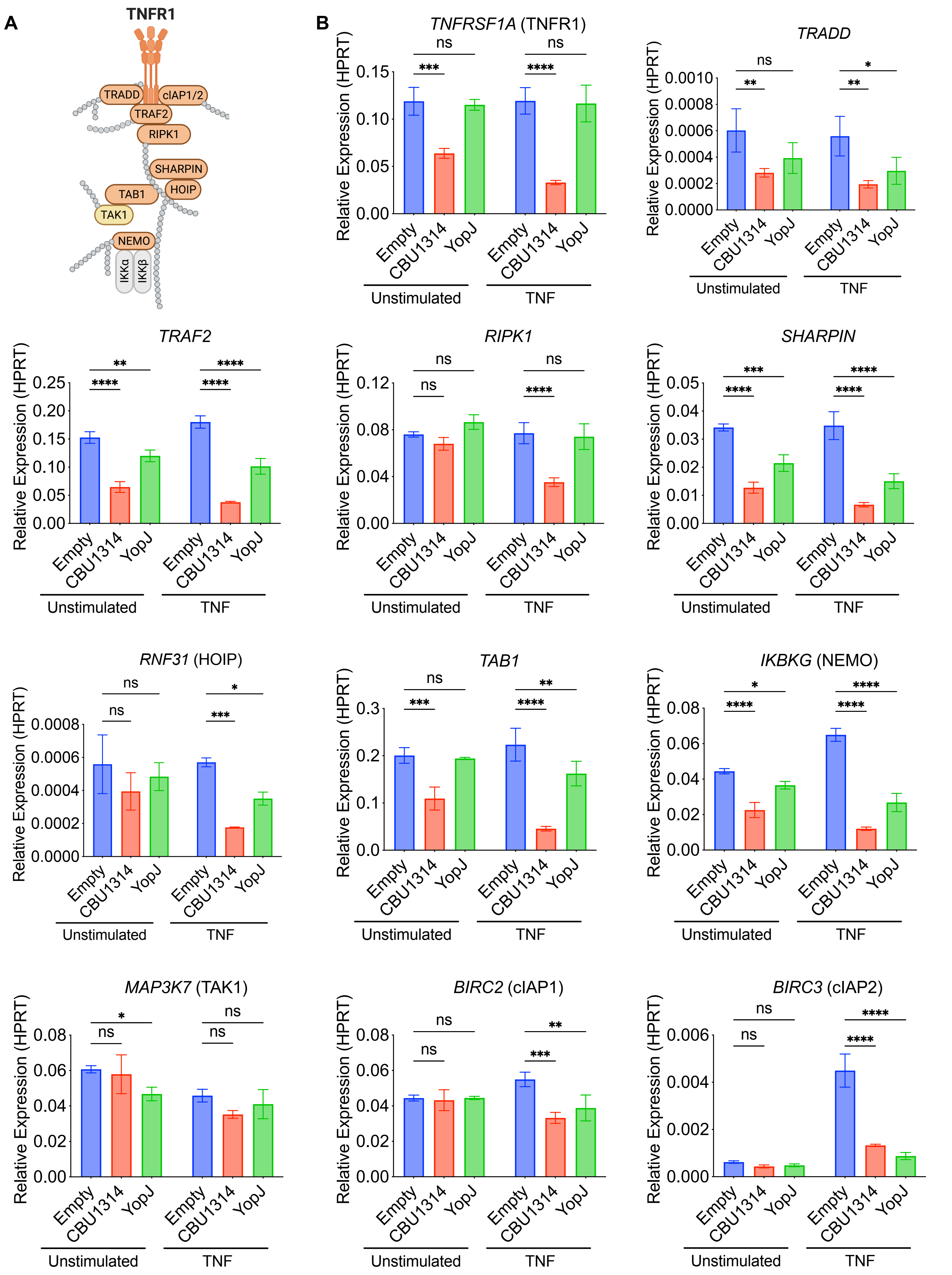

**Supplemental Figure 5. CBU1314 expression leads to decrease in mRNA levels of various TNFR1 pathway genes.** HEK293T cells were transfected with an empty vector or vectors encoding CBU1314 or YopJ. 24hrs later, cells were left unstimulated or stimulated with TNF for 6hrs prior to RNA harvest. A) mRNA expression of TNF pathway genes downstream of TNFR1 were analyzed. Orange indicates proteins with genes that were downregulated (B) and yellow indicates proteins with genes that were not affected (C) by CBU1314 expression. Gray indicates proteins that were not examined for gene expression. Graphical representation of TNFR1 pathway created with BioRender.com. Statistical analysis: Two-way ANOVA with Dunnett's multiple comparisons test within each unstimulated or TNF-stimulated group.

Supplemental Figure 6

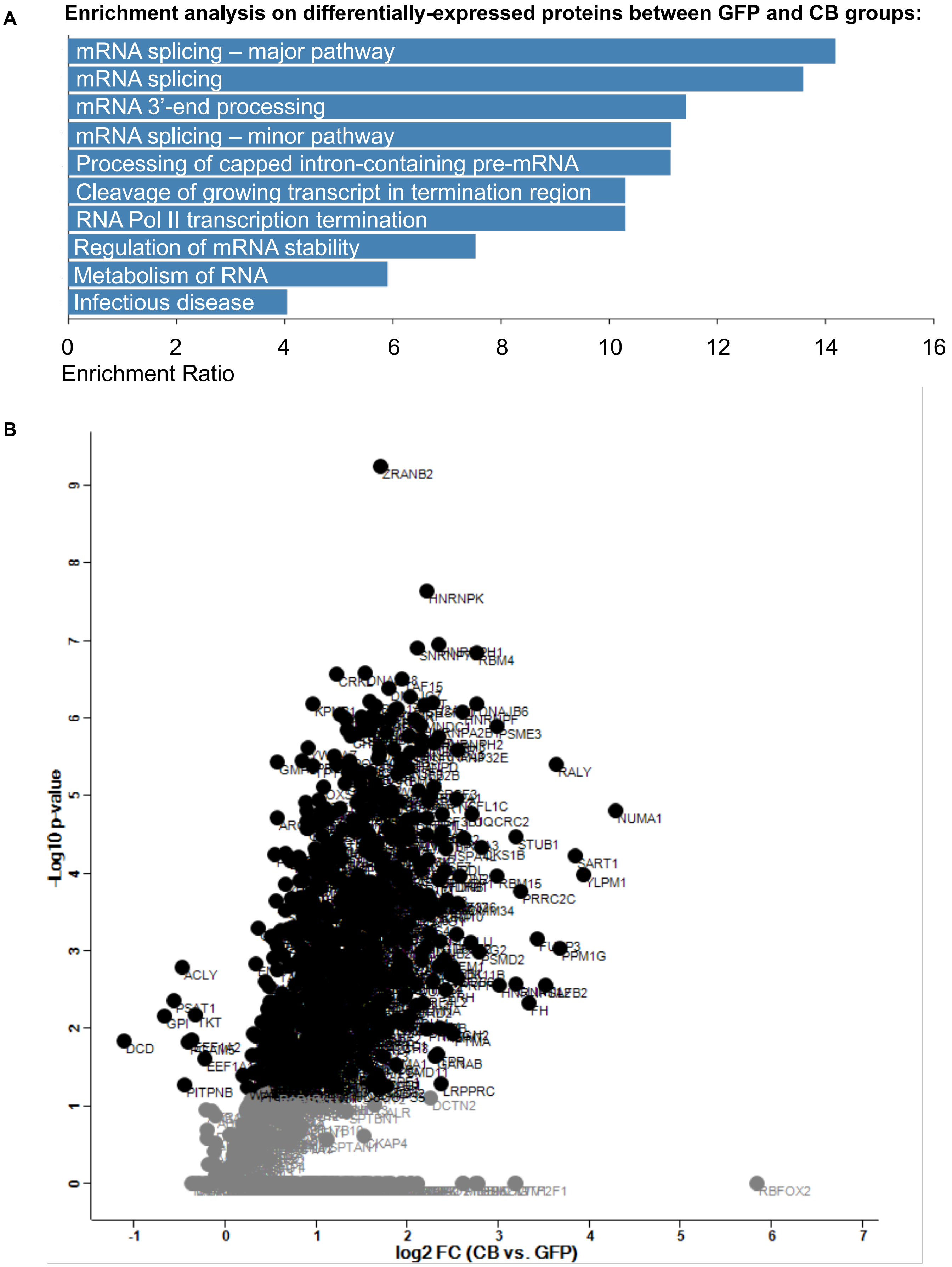

**Supplemental Figure 6. Pathway enrichment of differentially expressed proteins in the control and CBU1314 IP-MS samples**. A) A t-test analysis between the control GFP and experimental GFP-CBU1314 IP-MS samples was carried out to identify differentially expressed proteins. 504 proteins were identified as being significantly different between the groups, and pathway enrichment analysis was carried out in this group. B) Volcano plot displaying differentially expressed proteins between control (GFP) and experimental CBU1314 group (CB). Black dots indicate significant hits with $p<0.05$, while grey dots indicate insignificant hits, as determined by Student's t-test analysis.

### Supplemental Figure 7

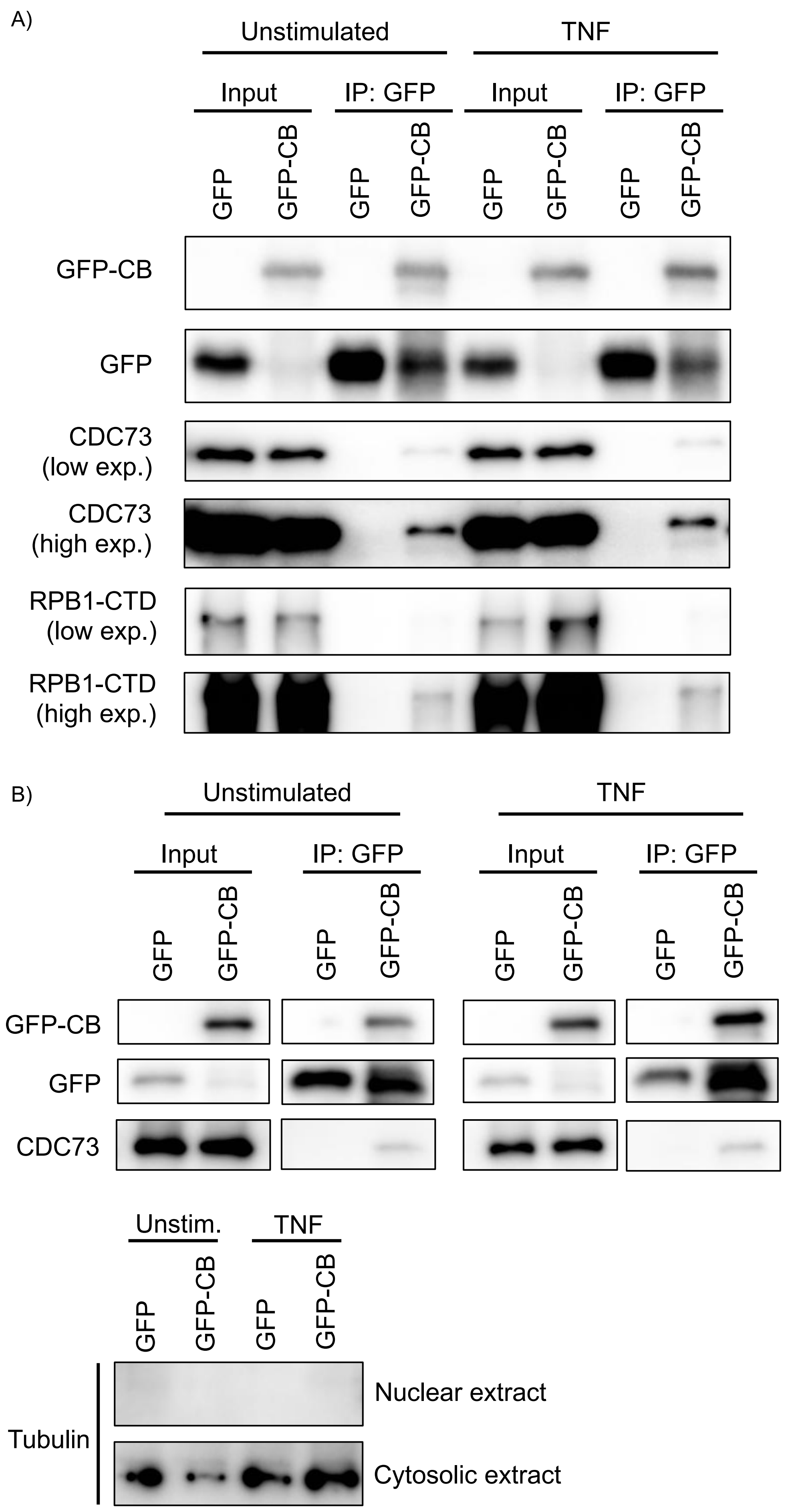

**Supplemental Fig. 7. CBU1314 interacts with PAF1C proteins in the nucleus independently of TNF signaling.** Co-IPs performed in either whole cell extracts (A) or nuclear extracts (B) of HEK293T cells expressing either GFP alone or GFP-CBU1314, and left unstimulated or stimulated with TNF. Immunoblot analysis using antibodies specific for GFP, CDC73, and RPB1-CTD. For B, immunoblotted for tubulin on input samples to verify purity of nuclear extracts.

### Supplemental Figure 8

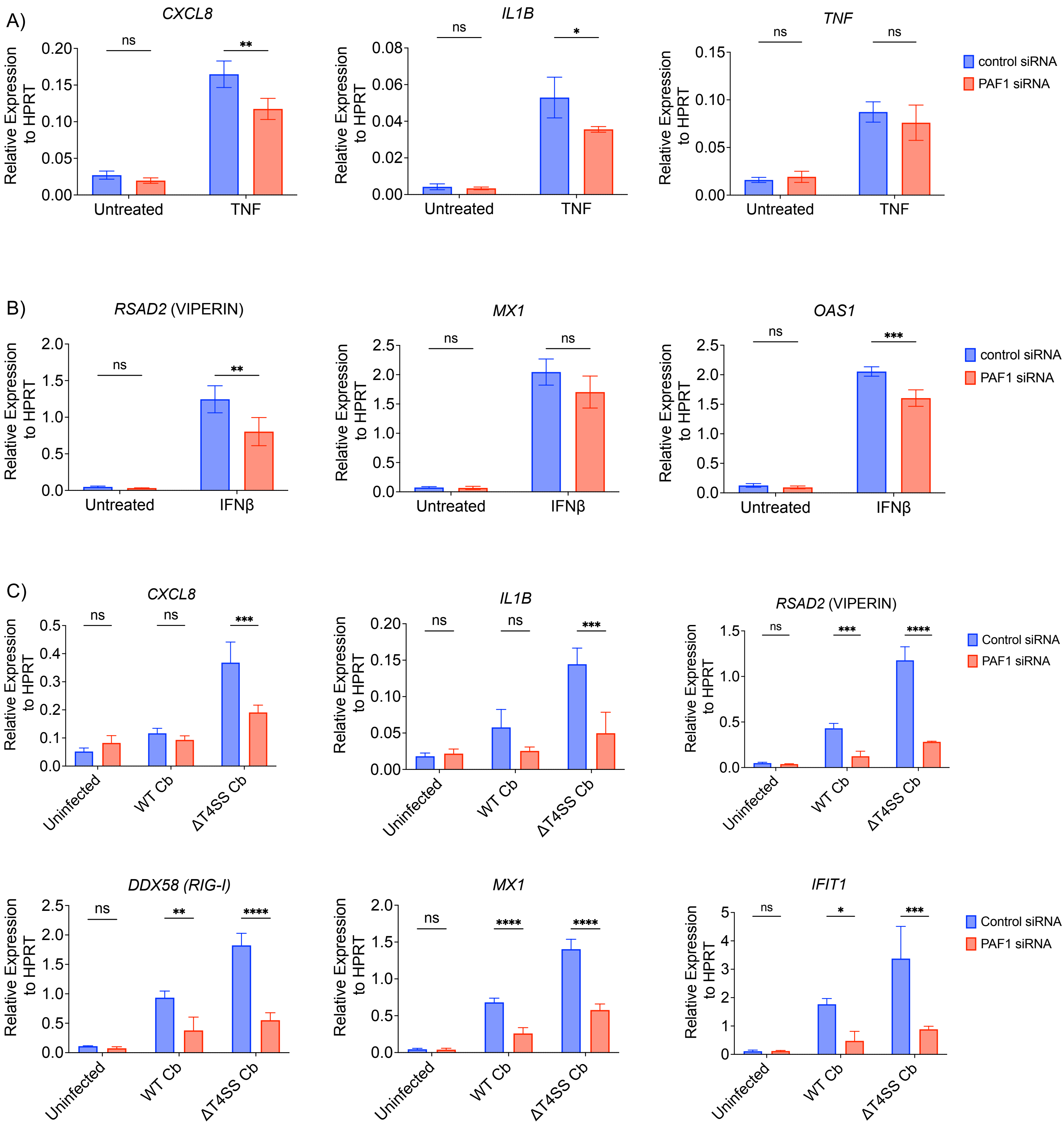

**Supplemental Figure 8. PAF1 contributes to gene expression downstream of TNF and IFN signaling and during infection.** THP-1 monocytes were treated with control or *PAF1* siRNA for 48hrs prior to stimulation with A) TNF, B) IFNβ or C) infection with the indicated *C. burnetii* strains (Cb). Samples were harvested for RNA analysis 6hrs post-stimulation, or 24hrs post-infection. Statistical analysis: Two-way ANOVA with Sidak's multiple comparisons test within each unstimulated or stimulated group, or infection condition.
